# Recursive feedback between Piezo1 conformation and membrane mechanics drives self-organization into finite clusters

**DOI:** 10.64898/2026.09.14.751624

**Authors:** Zixian Guo, Amrit Bagchi, Monika Dhankhar, Mohammad Dehghany, Vivek B. Shenoy

## Abstract

Piezo1 is a major mechanosensitive ion channel through which cells convert physical force into calcium-dependent signaling programs. In living membranes, this conversion depends not only on channel activation, but also on whether Piezo1 channels remain dispersed, assemble into finite clusters, or concentrate at sites where receptor signaling and mechanical forces reorganize the membrane. How single-channel force sensing is amplified into these collective spatial states remains unknown. Here we identify a membrane-feedback mechanism that converts single-channel mechanosensing into self-organized Piezo1 clusters. Coupling channel shape to membrane-cortex elasticity reveals that neighboring channels relax shared deformation fields, generating an effective interaction with short-range attraction opposed by longer-range repulsion. As channel density or membrane tension increases, this balanced interaction shifts Piezo1 from dispersed channels into mesoscale finite clusters. Brownian-dynamics simulations reproduce experimentally observed Piezo1 cluster geometries and swelling-induced cluster growth, while comparisons across distinct cellular systems place Piezo1 organization within a common density-tension framework. Applying the same mechanism to LPS-activated macrophages shows how receptor-induced membrane reorganization locally concentrates Piezo1 above the clustering threshold. Overall, these results recast Piezo1 mechanotransduction from isolated-channel force sensing to a membrane-driven self-organization process that spatially biases force-dependent calcium signaling within cells.

## Introduction

Piezo1 transforms mechanical forces at the cell surface into calcium signals that shape cellular behavior^1–3^. Through this function, Piezo1 regulates diverse physiological processes, including vascular development, epithelial homeostasis, immune function, red blood cell volume control, and cell fate decisions^1,4–16^. Most biophysical descriptions of Piezo1 have focused on how membrane tension promotes the opening of individual channels^4–9,17–23^. However, Piezo1 function in living membranes also depends on where channels are positioned^20^. Piezo1 channels can remain dispersed, assemble into finite clusters, or concentrate at sites where receptor signaling, adhesion, membrane curvature, or cortical mechanics reorganize the plasma membrane^4–9,17–23^. These distinct spatial states may determine whether mechanical inputs are sensed broadly across the cell surface or concentrated into localized zones of force-dependent calcium signaling. What determines these distinct spatial organizations and how they emerge from the mechanics of individual channels remain unclear.

A key challenge is that Piezo1 is not a passive membrane inclusion. Its large trimeric architecture deforms the surrounding lipid bilayer, while membrane tension and other mechanical inputs can shift the channel toward flatter shapes that favor opening and ion conductivity^2,3,24–29^. A Piezo1 channel that changes shape under force also changes the membrane deformation it imposes, and that altered deformation can influence nearby channels in its local membrane environment. This reciprocal coupling suggests that Piezo1 spatial organization cannot be understood from isolated-channel gating alone. How this single-channel force sensitivity is amplified into many-channel organization across the plasma membrane remains unknown. Existing theoretical and computational studies^21,28,30^ explain important components of Piezo1 mechanobiology, but leave unresolved how Piezo1 spatial organization emerges. Models of mechanosensitive-channel gating describe how membrane tension regulates isolated-channel opening, whereas structural and molecular simulations quantify Piezo1 deformation and force-induced flattening^2,3,21,24–27^. Separately, membrane-elasticity theories show how fixed-shape inclusions interact through deformation fields^31–37^. However, Piezo1 introduces feedback between channel shape and membrane mechanics, in which membrane tension changes its conformation and that conformational change alters the membrane deformation transmitted to neighboring channels. Existing approaches therefore miss the reciprocal feedback between channel state and membrane shape that is needed to predict whether Piezo1 remains dispersed, forms finite clusters, grows under osmotic swelling, or concentrates near receptor-associated membrane regions.

Here we develop a self-consistent membrane-feedback theory that links Piezo1 single-channel mechanics to collective membrane organization. In this framework, Piezo1 shape and deformation of the surrounding membrane-cortex composite are solved together, so that force-induced channel flattening determines how strongly each channel reshapes the membrane and interacts with its neighbors. We show that this reciprocal coupling generates a non-monotonic Piezo1-Piezo1 interaction that stabilizes finite clusters. The framework further predicts how channel density and membrane tension jointly control the transition from dispersed to clustered states. Brownian-dynamics simulations and comparisons across cellular systems support these predictions and show how mechanical perturbations reshape Piezo1 organization. Extending the same mechanism to spatially heterogeneous membranes further shows how local membrane remodeling can recruit Piezo1 and promote clustering at specific membrane regions. Together, these findings reveal how force sensing by individual Piezo1 channels can be amplified into spatially organized mechanotransduction across the cell membrane.

## Results

### Effective membrane-cortex mechanics determine the Piezo1 flattening state under tension

Piezo1 is a strongly curved mechanosensitive channel whose dome-like architecture deforms the surrounding cellular membrane (Fig. 1a). In the cell-based systems considered here^1,17,19,30^, this deformation is resisted not only by the lipid bilayer but also by the mechanically coupled cortical environment^38^. We therefore describe the local mechanical response using effective membrane-cortex properties (Fig. 1a). At the channel edge, the membrane meets Piezo1 at a finite angle, so changes in channel shape also change the deformation imposed on the surrounding bilayer (Fig. 1a). Motivated by structural and biophysical studies showing force-dependent flattening of Piezo1^27,39,40^, we parameterize this shape change with a membrane-coupled flattening coordinate *q*, ranging from a more curved, closed-like state at smaller values (*q* ≈ 0) to a flatter, more open-like state at larger values (*q* ≥ *q*_sat_) (Fig. 1a,b). Most membrane-mediated interaction models fix the channel flattening state first and then compute the deformation around that prescribed state^32–34,41^. Piezo1, however, requires a different description because membrane tension can shift the channel flattening state itself. Increasing *q* expands the projected channel footprint and reduces the boundary mismatch imposed on the surrounding membrane. We therefore first determine how membrane tension shifts the Piezo1 flattening state and how much membrane deformation remains after the channel and the surrounding membrane-cortex composite relax together. This remaining deformation provides the mechanical source through which Piezo1 later couples to nearby channels and localized membrane structures.

The Piezo1 flattening state is set by a balance between channel elasticity and the mechanical response of the surrounding cellular membrane^39,40^. Three contributions are central: the protein’s intrinsic preference for its reference conformation, the mechanical work gained when the projected footprint expands under membrane tension, and the elastic cost of matching the surrounding membrane-cortex composite to the channel edge (Fig. 1c-e). We therefore write the single-channel free energy as the sum of a channel contribution and an effective membrane-mechanical contribution, denoted by subscripts ch and mem, respectively. In this coarse-grained description, the membrane deformation is represented by a height field *u*(**x**), with *κ*_eff_ denoting the effective bending modulus of the membrane-cortex composite and *σ* representing the effective membrane tension transmitted through the membrane-cortex system and acting on the channel. The total single-channel free energy can be written as

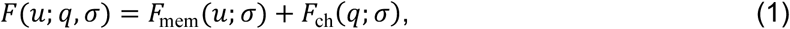

with

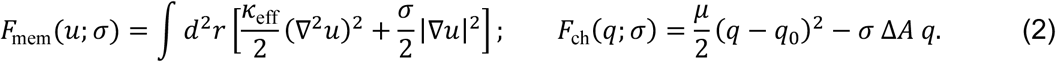

Here *μ* is the stiffness of the flattening coordinate, *q*_0_ is the preferred state of the protein in the absence of membrane forcing, and *ΔA* is the projected area gain per unit increase in *q* (Fig. 1e). Mechanically, membrane tension can shift Piezo1 away from its intrinsic state because increasing *q* costs conformational energy but gains mechanical work through footprint expansion and can reduce the deformation that must be imposed at the channel edge.

The coupling between Piezo1 shape and cellular membrane deformation enters through the boundary condition at the channel edge. In the small-slope limit, the edge tilt is the normal slope of the membrane height field^41^, so we impose *∂_n_u*|_edge_ = *α*(*q*) at the channel boundary, with the deformation decaying back to the far-field membrane state away from the channel. More curved Piezo1 states impose a larger edge tilt *α*(*q*), whereas flatter states impose a smaller one (Fig. 1a,b). We therefore write the state-dependent boundary angle as *α*(*q*) = [*α_c_* − *λq*]_+_, where [*x*]_+_ ≡ max(*x*, 0). Here, *α_c_* is the edge tilt of the most curved state, and feedback strength *λ* measures how efficiently channel flattening reduces the boundary mismatch imposed on the surrounding membrane-cortex composite (Fig. 1b). The positive part operator means that once Piezo1 has flattened enough to remove the imposed edge tilt, further increases in *q* no longer change the membrane boundary condition. This saturation occurs at *q*_sat_ = *α_c_* / *λ*. The resulting tension-dependent progression toward this saturated state, together with the corresponding reduction in residual boundary mismatch, is shown in Fig. 1b and Supplementary Fig. 1a,b.

**Figure 1:**
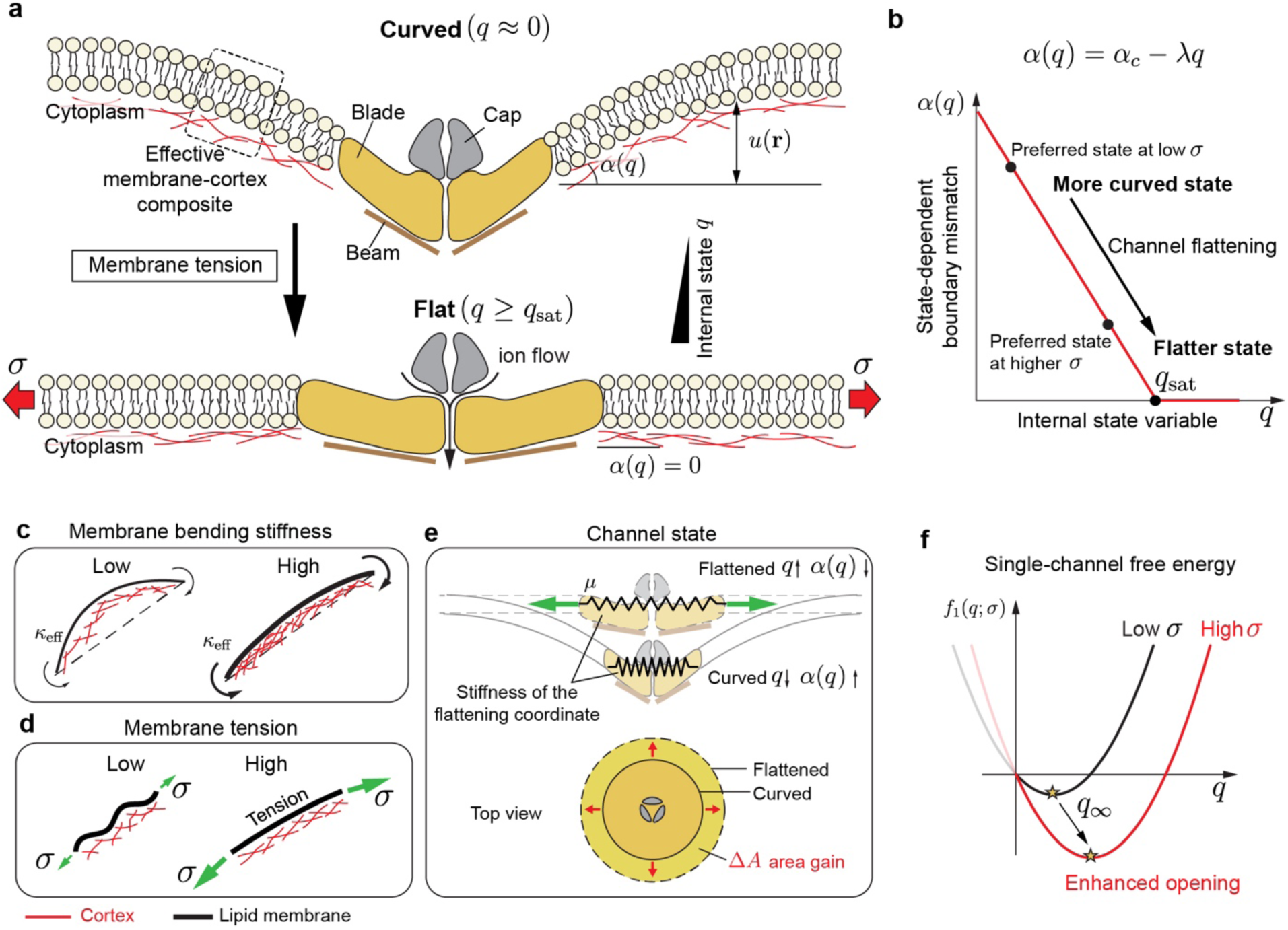
Single-channel mechanics couple Piezo1 conformation to the local membrane deformation field. **a,** Schematic of Piezo1 embedded in the membrane in more curved, closed-like and flatter, open-like conformations. Channel flattening changes the boundary mismatch imposed on the surrounding membrane and is promoted by membrane tension. **b,** State-dependent boundary mismatch, written as *α*(*q*) = *α_c_* − *λq* on the active branch, decreases with the internal state variable *q* and saturates at *q*_sat_. Stronger feedback *λ* produces a steeper reduction in mismatch with flattening. **c,** Effective membrane-cortex stiffness as a coarse-grained input to the one-body mechanics. **d,** Membrane tension as the mechanical input that biases channel flattening and opening. **e,** Schematic of the state-dependent membrane footprint of Piezo1. Flattening toward the open state reduces the imposed boundary mismatch and increases the projected membrane area. **f,** Reduced single-channel free energy *f*_1_(*q*; *σ*) illustrating the shift of the preferred state toward larger *q* at higher membrane tension. The relaxed one-body outputs, including the isolated-channel state *q*_∞_ and residual mismatch *α*_∞_, provide the inputs to the pair-interaction theory developed below.

For any chosen channel state *q*, the imposed edge tilt *α*(*q*) determines how Piezo1 mechanically loads the surrounding cellular membrane. The membrane-cortex composite then relaxes around this boundary condition and returns to the far-field state away from the channel. In Supplementary Note 1, we solve this height-field minimization explicitly and reduce the membrane contribution to an effective cost of imposing the remaining edge tilt on the surrounding membrane. Substituting the relaxed deformation back into the energy gives a reduced single-channel free energy that depends only on the channel state and this remaining edge tilt. The resulting single-channel free energy is

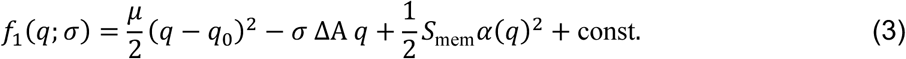

Here, *S*_mem_ represents the effective stiffness of the surrounding membrane-cortex composite against the edge tilt imposed by Piezo1. As derived in Supplementary Note 1, *S*_mem_ is obtained by minimizing the membrane deformation for a prescribed boundary slope, reducing the distributed membrane elastic energy to the effective cost *S*_mem_*α*(*q*)^2^/2. A larger *S*_mem_ therefore makes the same residual edge tilt more energetically costly. Equation (3) then captures the competition between the intrinsic preference of Piezo1 for its reference state, the work gained by footprint expansion under membrane tension, and the membrane elastic cost of the remaining edge tilt (Supplementary Fig. 1c). We minimize the reduced free energy with respect to *q* to determine the relaxed state of an isolated channel.

The resulting relaxed isolated-channel state *q*_∞_(*σ*) is the flattening state approached by a channel in an otherwise uniform membrane, far from other channels or localized membrane structures (hence the subscript ∞). Minimizing the reduced free energy with respect to the flattening coordinate gives

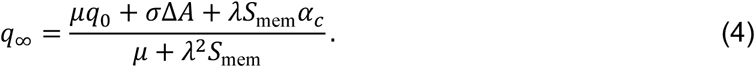

This expression shows how membrane tension shifts the channel away from its intrinsic state. The numerator contains the three drives toward a larger flattening state: the intrinsic bias *μq*_0_, the tension work *σ*Δ*A*, and the elastic gain in the membrane-cortex system from reducing the Piezo1-imposed edge tilt. The denominator is the total stiffness resisting changes in *q*, combining the intrinsic channel stiffness with the stiffness of the surrounding membrane-cortex composite. This result shows that Piezo1 does not impose a fixed edge tilt on its surroundings. Instead, the edge tilt that remains after channel relaxation depends on the local membrane tension and on the elastic properties of the membrane-cortex composite. Higher tension shifts Piezo1 toward a flatter state, whereas a stiffer membrane-cortex composite enhances the elastic cost of the edge tilt that the channel imposes (Fig. 1f). The relaxed state *q*_∞_ therefore determines both how open-like the isolated channel becomes and how strongly it loads its mechanical environment. These tension- and stiffness-dependent changes in channel shape and edge tilt provide the mechanical input for Piezo1-Piezo1 interactions in the next section.

### Recursive state-shape feedback generates short-range attraction and long-range repulsion between Piezo1 channels

We next asked how the tension- and stiffness-dependent edge tilt derived for an isolated Piezo1 channel changes when two channels are close enough to deform the same membrane-cortex composite. The single-channel calculation shows that each Piezo1 molecule is characterized by a relaxed state *q*_∞_(*σ*), and by the remaining edge tilt that it imposes on its surroundings. In a two-channel system, these quantities become coupled such that the deformation generated by one channel changes the local membrane shape sensed by the other, shifting the preferred state of both channels toward a flatter conformation (Fig. 2a-c). This neighbor-induced flattening reduces the edge tilt that the second channel imposes back on the membrane-cortex composite and lowers the elastic energy stored in the shared deformation field (Fig. 2d). Piezo1-Piezo1 interactions therefore cannot be described as passive overlap between two fixed deformation fields. They are reshaped by the conformational response of each channel to the deformation generated by its neighbor.

**Figure 2:**
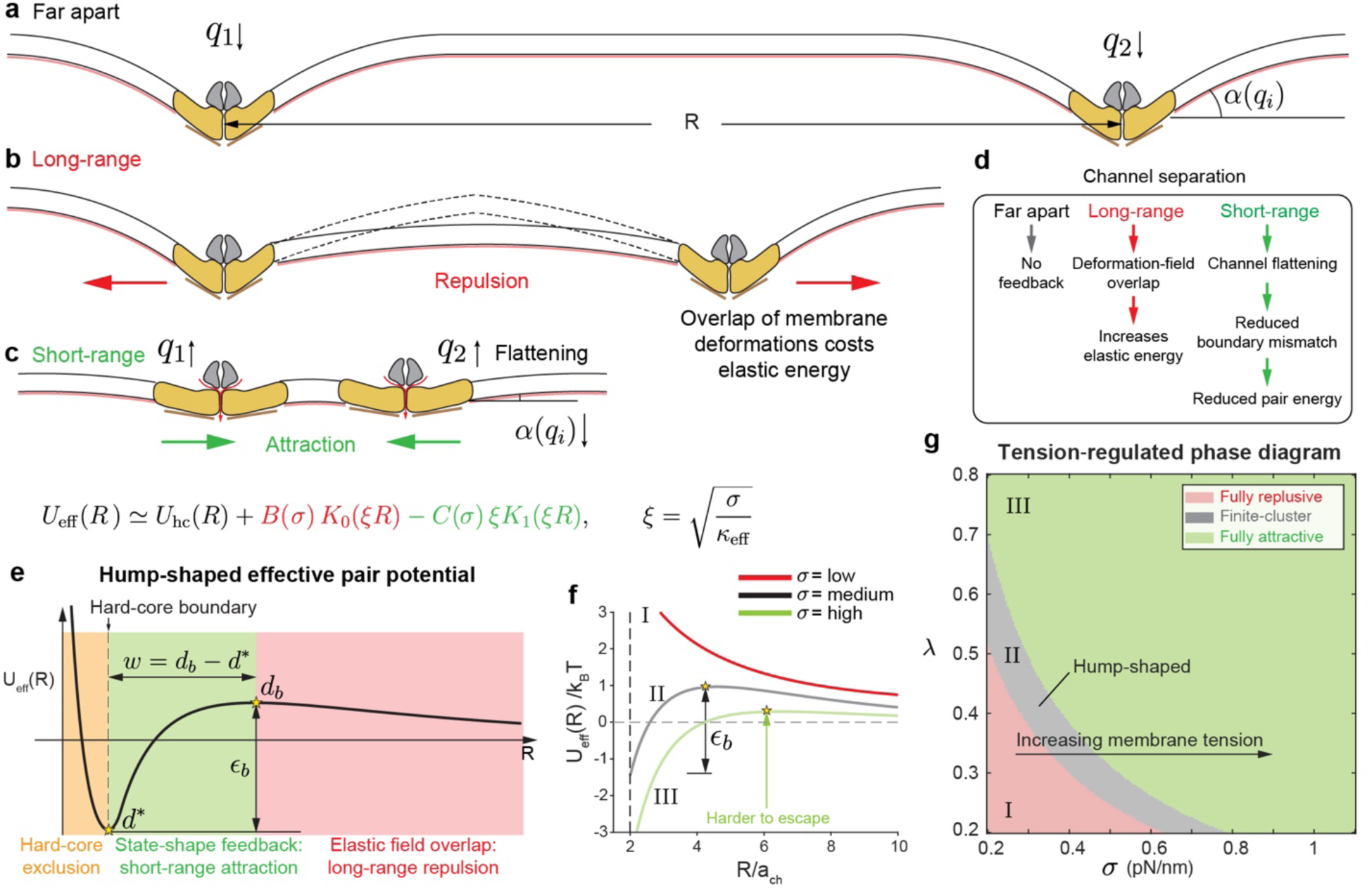
Recursive state-shape feedback generates a non-monotonic Piezo1 pair potential. **a**, Two isolated channels at large separation each generate an independent membrane deformation field. **b,** Direct overlap of the membrane deformation fields increases elastic energy and gives a longer-range repulsive contribution. **c,** At shorter separation, the deformation generated by one channel shifts the neighboring channel toward a flatter state, reducing its boundary mismatch and lowering the pair energy. **d,** Schematic summary of the recursive feedback pathway linking membrane deformation, channel flattening, reduced boundary mismatch, and reduced pair energy. **e,** Effective pair potential *U*_eff_(*R*) showing hard-core exclusion, a short-range attractive branch generated by state-shape feedback, and a longer-range repulsive tail arising from elastic-field overlap. The preferred separation *d*^∗^, outer barrier *d_b_*, attractive-basin width *w* = *d_b_* − *d*^∗^, and escape barrier *ε_b_* are indicated. **f,** Representative pair potentials at different membrane tensions. Increasing tension deepens the attractive well and lowers the barrier to pair association within the finite-cluster regime. (The dashed line represents the hard-core boundary) **g,** Tension-regulated phase diagram in the (*λ*, *σ*) plane showing fully repulsive regime, hump-shaped finite-cluster regime, and fully attractive regime.

To quantify this coupling, we compute the two-channel free energy for channels separated by a center-to-center distance *R*. For each separation, the membrane-cortex deformation is relaxed subject to the edge tilts imposed by both channels, and the channel states are expanded about the isolated-channel state *q*_∞_(*σ*) derived above. Eliminating the shared membrane deformation field and expanding the channel state about the isolated-channel solution, as derived in Supplementary Note 2, gives an effective pair potential:

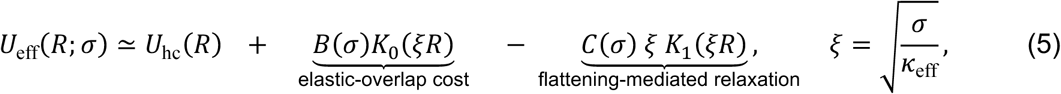

where *U*_hc_(*R*) enforces steric hard-core exclusion between channel footprints with *d*^∗^ denoting the minimum allowed center-to-center separation, set by the physical size of the Piezo1 membrane footprint, below which two channels cannot overlap (Fig. 2e). The remaining terms describe membrane-mediated interactions beyond this hard-core contact distance. The functions *K*_0_ and *K*_1_ are modified Bessel functions of the second kind, and 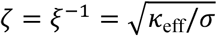 represents the characteristic membrane screening length. This length sets the distance over which a Piezo1-induced membrane deformation persists before being damped by tension. Thus, increasing membrane tension shortens the interaction range and reshapes the pair potential. The interaction amplitudes are governed directly by the isolated-channel mechanical properties, defined as 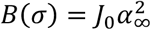 and 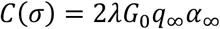 . Here, the constant 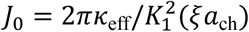 sets the direct elastic cost of field overlap, where *a*_ch_ is the effective Piezo1 channel footprint radius and *α*_∞_ = *α_c_* − *λq*_∞_ is the relaxed isolated-channel boundary mismatch. The quantity *G*_0_ = *a*_ch_*J*_0_ then parameterizes the magnitude of the neighbor-induced shift in the conformational state obtained in the pair-state expansion derived in Supplementary Note 2. Physically, the coefficient *B* quantifies the elastic penalty of overlapping the residual membrane distortions produced by two incompletely relaxed channels. When these deformation fields overlap, they increase local curvature gradients and raise the membrane elastic energy. In contrast, the coefficient *C* quantifies the elastic energy saved when one channel’s deformation field drives its neighbor toward a flatter conformation that imposes a smaller boundary tilt on the surrounding bilayer. The opposite signs of these two terms therefore encode a direct physical competition: deformation-field overlap is repulsive, whereas neighbor-assisted reduction of the imposed boundary slope is attractive. Importantly, this attraction is not a phenomenologically added interaction but emerges intrinsically from the state-dependent boundary condition of Piezo1 itself.

Because deformation-field overlap and neighbor-assisted flattening decay differently with separation, their competition produces a non-monotonic pair potential rather than a uniformly attractive or uniformly repulsive interaction (Fig. 2e). At separation below the hard-core distance *d*^∗^, steric exclusion prevents overlap of the Piezo1 footprints. Immediately beyond this contact distance, the neighbor-assisted flattening term begins to dominate. Each channel can reduce the boundary tilt imposed on the bilayer by responding to the deformation field of the other channel. This relaxation lowers the membrane elastic energy and produces a short-range attractive well. At larger separations, the conformational response decays more rapidly, whereas the residual deformation fields still overlap over the membrane screening length. The remaining interaction is therefore dominated by the elastic cost of field overlap, producing an outer repulsive barrier (Fig. 2e).

The shape of the Piezo1-Piezo1 potential defines the physical length and energy scales that control pair association. The minimum of the attractive basin defines the preferred pair spacing, while the outer maximum *d_b_* (satisfying 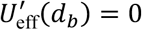 defines the repulsive barrier that a pair must cross to dissociate from the attractive basin (Fig. 2e). The basin width, *w* = *d_b_* − *d*^∗^, measures the range of separations over which two channels are effectively captured, and the energy difference between the well minimum and the outer barrier defines the escape barrier, 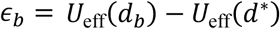 . These quantities reduce the continuous pair potential to a small set of spacing, capture, and escape scales that can be used directly in the theory to predict channel clustering.

The strength of conformational feedback *λ* determines whether the attractive basin exists at all (Supplementary Fig. 2). If flattening only weakly reduces the boundary tilt imposed by Piezo1, the relaxation term is too small to overcome the elastic penalty of overlapping residual membrane deformations, and the interaction remains effectively repulsive outside the hard-core contact distance (Supplementary Fig. 3). As the feedback strength increases, neighbor-assisted flattening becomes more effective, the attractive well deepens, the preferred spacing shifts closer to contact, and the escape barrier increases. In this regime, two channels that enter the attractive basin are more likely to remain associated. When the feedback is suppressed, the attractive branch disappears and the pair potential collapses to the purely repulsive limit (Supplementary Fig. 3).

Membrane tension tunes this same interaction through two coupled effects. First, increasing tension shifts the isolated-channel free-energy minimum toward a flatter state *q*_∞_, thereby reducing the edge tilt that the neighboring channel imposes on the membrane-cortex composite and changing the amplitudes of both the repulsive and attractive terms. Second, increasing tension shortens the membrane screening length 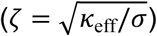, limiting the distance over which Piezo1-induced deformation propagates. Tension therefore does not simply strengthen or weaken the interaction but reshapes the potential topology by simultaneously changing the channel conformation and the spatial decay of the membrane deformation field. Within the parameter range examined here, higher tension lowers the outer repulsive barrier and makes the attractive basin more accessible (Fig. 2f).

Together, these effects define three regimes of the pair potential (Fig. 2g), which we categorize as fully repulsive, finite-cluster, and fully attractive. Because membrane tension and conformational feedback control different parts of the interaction, the boundaries between these regimes depend on both variables. Feedback sets how effectively neighbor-induced deformation flattens a channel and reduces its imposed edge tilt, whereas membrane tension shifts the isolated-channel state and changes the screening length over which deformation propagates (Fig. 2g). In the repulsive regime, channel relaxation is too weak to overcome the elastic cost of deformation-field overlap. In the finite-cluster regime, short-range attraction coexists with a longer-range repulsive barrier, producing a preferred spacing, a finite capture basin, and a nonzero escape barrier. In the fully attractive regime, the repulsive barrier is lost and the pair potential no longer contains an intrinsic outer length scale that limits pair association. Thus, recursive state-shape coupling generates the non-monotonic interaction needed for finite Piezo1 clustering only within a defined window of membrane tension and feedback strength.

### Density and membrane tension determine the onset of finite Piezo1 clustering

We next asked how the pair interaction derived above gives rise to collective Piezo1 organization at the mesoscopic scale. The model predicts a progression from dispersed channels to small oligomers and finite nanoscale assemblies as illustrated by the representative configurations in Fig. 3a,b. These distinct spatial states arise from two physical ingredients in our framework: enough channels for repeated encounters, and a membrane-cortex mechanical state that makes those encounters energetically favorable. Channel density controls how frequently diffusing Piezo1 molecules enter the short-range attractive basin of *U*_eff_(*R*), whereas membrane tension controls the shape of that basin by changing the isolated-channel state, the remaining edge tilt, and the screening length (Fig. 3b). At low areal density, encounters are rare, so the membrane is populated mainly by dispersed monomers and transient small oligomers. As density increases, channels enter the attractive basin more often, allowing dimers and larger assemblies to become stabilized by the pair interaction. At the same time, the longer-range repulsive branch penalizes continued growth and prevents collapse into one macroscopic aggregate. The result is a density- and tension-dependent transition from dispersed Piezo1 channels to stable finite clusters.

This transition can be expressed quantitatively using the same geometric and energetic features of the pair potential that control two-channel association. The relevant length scale is the capture area associated with the attractive basin *A_b_*(*σ*, *λ*), which measures the area around a channel within which another channel can be trapped by the short-range attraction (Fig. 3a). The relevant energetic scale is the escape barrier *ε_b_*(*σ*, *λ*), which measures how much thermal energy is required for an associated pair to leave that basin. Defining 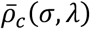 as the mean areal channel density and *β* as the inverse thermal energy, the onset of finite clustering occurs when

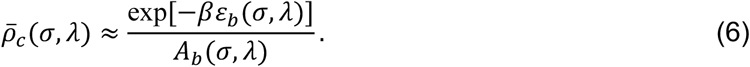

This condition states that clustering begins when channels both find the attractive basin often enough and remain there long enough to stabilize pairs. The capture area sets the likelihood that a diffusing channel encounters a neighbor within the attractive range, while the escape barrier sets the thermal stability of that captured pair. A larger capture area increases the probability of pair formation, and a larger escape barrier makes pair dissociation less likely. Both effects lower the channel density required to transition from the dispersed regime into the finite-cluster regime. Because both quantities are determined by the shape of *U*_eff_(*R*), which varies with membrane tension *σ* and feedback strength *λ*, this onset condition also defines a phase diagram for the predicted onset channel density *ρ_c_*(*σ*, *λ*)(Supplementary Fig. 4b). Supplementary Note 3 derives this onset criterion from the expected number of neighboring channels thermally captured within the attractive basin.

### Brownian-dynamics simulations connect pair interactions to finite Piezo1 organization

To test whether this pair-level onset criterion predicts collective organization in many-channel systems, we performed Brownian-dynamics simulations in which Piezo1 channels diffuse under thermal fluctuations and forces from the derived pair potential. This allowed us to ask whether the spacing, capture, and escape scales obtained from the two-channel calculation are sufficient to generate finite clusters in a fluctuating many-channel membrane. At low channel density, channels rarely enter the attractive basin and remain mostly dispersed, with only transient dimers and small oligomers (Fig. 3b,c). As density increases, repeated entry into the attractive basin stabilizes longer-lived finite assemblies (Fig. 3b,c). Channels within each assembly move collectively, causing the center-of-mass diffusivity to decrease with cluster size because the drag contributions of the constituent channels accumulate. Accordingly, the simulated diffusivity *D_N_* of an N-channel cluster decreases approximately as *D_N_*/*D*_1_∼*N*^−1^, where *D*_1_ is the monomer diffusivity (Fig. 3d). Consistent with the analytical onset landscape in Supplementary Fig. 4b, the many-body clustered fraction in Supplementary Fig. 5 shifts to lower density at higher membrane tension. Higher tension reshapes the pair potential by making the attractive basin more accessible, thereby shifting the system more readily from dispersed channels into the finite-cluster regime (Fig. 2f).

The simulated spatial configurations illustrate the combined effects of channel density and membrane tension (Fig. 3e). At low density, channels remain mostly dispersed under both lower- and higher-tension conditions. At intermediate density, higher membrane tension shifts the system into discrete finite assemblies, whereas lower tension still favors monomers and small oligomers. At high density, clustering occurs under both tension conditions, but higher tension produces larger and more prevalent assemblies. These configurations provide the spatial counterpart to the onset curves in Fig. 3c, showing how the same pair potential generates dispersed, intermediate, and clustered states as channel density and membrane tension are varied.

We next compared the predicted finite-cluster regime with Piezo1 organization in HEK293T cells stably expressing Piezo1-GFP. Published images^18^ from this system were segmented to identify Piezo1-positive clusters at the plasma membrane, providing two experimental quantities for model comparison: the areal density of detected Piezo1 channels and the resulting cluster geometry. The measured Piezo1 channel density was used to set the number of simulated channels per unit membrane area in the Brownian-dynamics simulations (Fig. 3f,g). The overall workflow linking experimentally measured density, simulated cluster organization, and membrane-context inference is summarized in Supplementary Fig. 6. The resulting simulated particle configurations were then rendered into confocal-like images and analyzed with the same geometric segmentation as the experimental data. Supplementary Fig. 7 illustrates this conversion from a Brownian-dynamics particle configuration to its corresponding diffraction-limited morphology. This allowed the predicted cluster-area and cluster-perimeter distributions to be compared directly with the HEK293T measurements (Fig. 3f-j). The intrinsic Piezo1 parameters used in these simulations, including the channel footprint radius, projected area gains during flattening, closed-state boundary mismatch, feedback strength, flattening stiffness, and zero-tension reference state, were fixed from the literature-based coarse-grained model. Their definitions, physical roles, and literature basis are summarized in Supplementary Note 4 and Supplementary Tables 1-3.

In this comparison, membrane tension was treated as the remaining mechanical control variable after fixing the experimentally measured Piezo1 density. Changing the tension changes the isolated-channel state, the residual boundary mismatch, and the membrane screening length, thereby reshaping the same pair potential used in the Brownian-dynamics simulations. We therefore varied membrane tension to identify the membrane context in which the simulated organization matched the HEK293T cluster scale. This comparison tests the model more stringently than matching a single mean cluster size because the same density, intrinsic Piezo1 parameters, and membrane tension must reproduce the experimentally measured distributions of both cluster area and perimeter. The Brownian-dynamics simulations reproduce the broad, right-skewed shapes of both distributions (Fig. 3i,j), with cluster area reporting the distribution of assembly size and perimeter providing additional sensitivity to cluster boundary geometry and compactness.

The asymmetric distributions in Fig. 3i,j provide a consistent readout of the underlying attraction-repulsion balance. In the area distribution, small clusters become likely once channels can enter the short-range attractive basin, producing a peak at finite area rather than a purely dispersed population (Fig. 3i). However, the probability of very large clusters falls off because continued growth accumulates long-range membrane-mediated repulsion and boundary penalties, suppressing macroscopic coarsening. The perimeter distribution reflects the same physics expressed through cluster boundary geometry. Compact clusters with moderate perimeters are favored, whereas very large or highly extended boundaries are progressively disfavored by the same finite-cluster free-energy cost. Supplementary Note 5 formalizes this picture by coarse graining the pair interaction into a cluster free energy, in which short-range cohesion favors growth, while the loss of favorable contacts at the cluster boundary and accumulated longer-range repulsion limit continued expansion. In the compact-cluster limit, the perimeter distribution follows from the same free energy through the geometric relation between cluster area and perimeter. Thus, the agreement between the simulated and experimental cluster-area and cluster-perimeter distributions provides a stringent test of the model, as the same non-monotonic pair interaction that predicts the density- and tension-dependent clustering onset also reproduces the finite, right-skewed organization of Piezo1 clusters observed in HEK293T cells. These results support the conclusion that Piezo1 clusters in HEK293T cells reflect a mechanically selected finite-cluster state, rather than a nonspecific accumulation of channels at the membrane.

**Figure 3:**
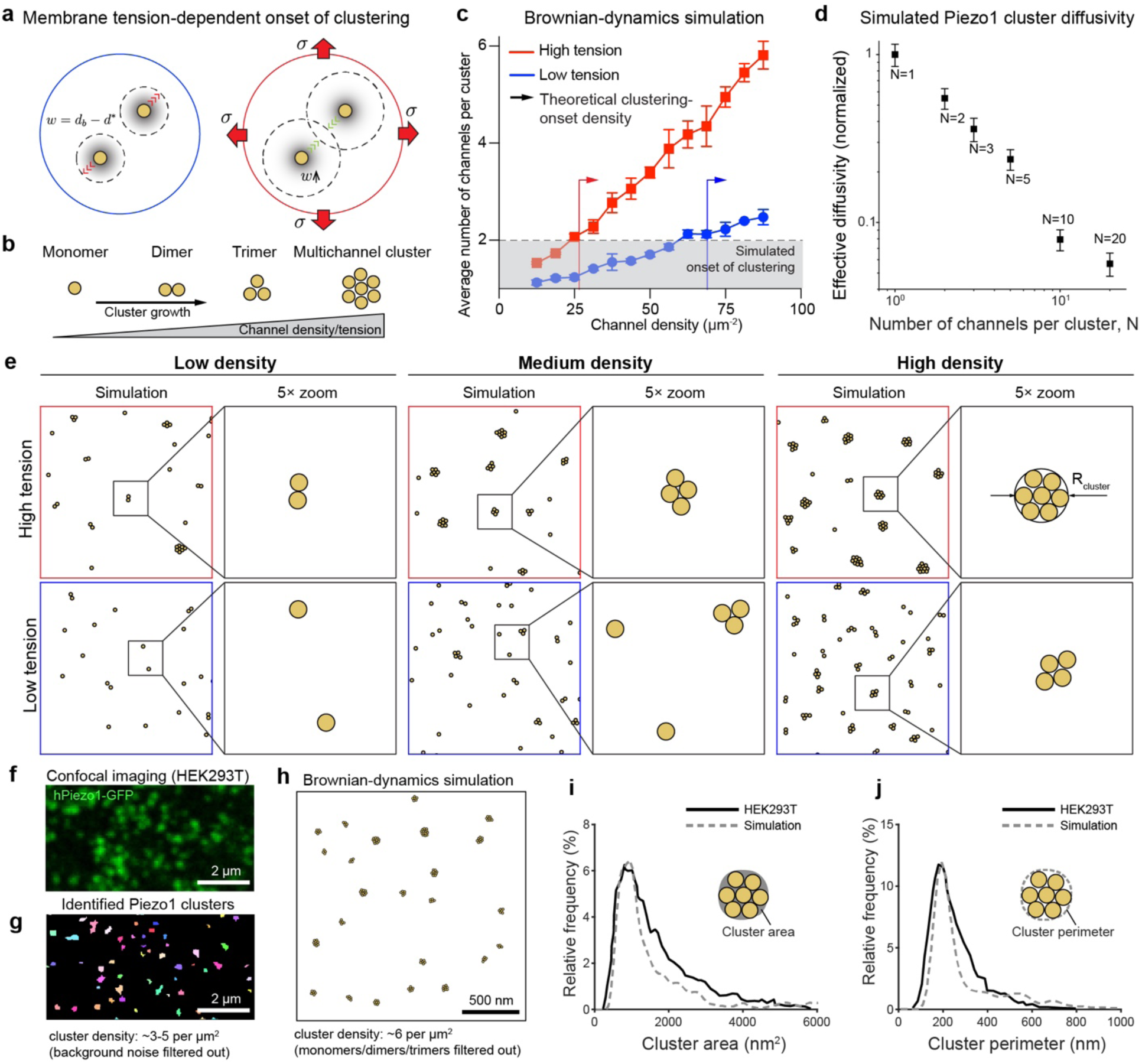
Density and membrane tension set the many-body states generated by the derived Piezo1 pair interaction. **a**, Schematic of membrane-tension-dependent channel trapping. Increasing membrane tension reshapes the pair interaction so that neighboring channels that encounter one another are more readily captured within the attractive basin. **b,** Representative progression from monomers to dimers, trimers, and multi-channel clusters. **c,** Mean number of channels per cluster as a function of channel density under high- and low-tension conditions. Higher membrane tension lowers the density required for clustering. Arrowed lines indicate the theoretical and simulated onsets of clustering. (n = 3; data are the mean ± s.d.) **d,** Normalized effective diffusivity as a function of number of channels per cluster. Brownian-dynamics simulations show that diffusivity decreases with cluster size, providing a kinetic mechanism that stabilizes larger finite assemblies. (n = 30; data are the mean ± s.d.) **e,** Brownian-dynamics configurations at low, intermediate, and high channel density under high- and low-tension conditions, shown together with magnified views of representative local configurations. High tension promotes finite cluster formation at lower density and increases cluster prevalence at matched density. **f,g,** Published confocal and cluster-segmented images of Piezo1-GFP in HEK293T cells, showing punctate nanoscale organization at the plasma membrane. **h,** Brownian-dynamics snapshot generated from the derived pair interaction at matched channel density for comparison with the HEK293T imaging phenotype. **i,j,** Cluster-area and cluster-perimeter distributions from HEK293T imaging and Brownian-dynamics simulation, with schematics indicating the geometric definitions of cluster area and cluster perimeter used in the analysis. The broad but finite distributions support the self-limited clustering regime generated by the model.

### Hypo-osmotic swelling reveals tension-driven Piezo1 cluster growth

We next asked whether an acute change in membrane mechanics is sufficient to move Piezo1 along the density-tension organization landscape predicted by the model (Fig. 3c,e). Hypo-osmotic swelling provides a direct perturbation of this axis: osmotic water influx expands the cell and increases membrane tension, while the intrinsic Piezo1 parameters remain unchanged on the timescale of the experiment (Fig. 4a). In the model, this increase in membrane tension shifts Piezo1 toward a flatter preferred conformation, reduces the residual boundary mismatch, and changes the membrane screening length, thereby reshaping the same non-monotonic pair potential that controls finite-cluster formation. Thus, using the experimentally measured Piezo1 density for each condition, the model predicts that hypo-osmotic swelling should shift Piezo1 populations toward larger and more cohesive finite assemblies by driving the membrane deeper into the clustered regime.

We tested this prediction using published Minimal Fluorescence Photon Fluxes (MINFLUX) localization data^17^ from Neuro-2a cells expressing Piezo1 under isotonic control and acute hypo-osmotic conditions, which provide nanometer-scale localization of individual fluorophores. The localization maps were converted into spatial density maps using an established Voronoi-tessellation pipeline developed in our earlier studies^42–44^, in which smaller Voronoi polygons indicate higher local Piezo1 density. The same segmentation parameters were applied to the isotonic and hypo-osmotic datasets, and contiguous high-density regions were grouped into clusters for geometric quantification. This analysis provided the experimental quantities needed for comparison with the model, including the relative Piezo1 density in each condition and the corresponding cluster-area distribution (Fig. 4b,c). Thus, the swelling experiment provides both the measured Piezo1 density and an independently defined mechanical perturbation, allowing the model to predict how Piezo1 organization should change under increased membrane tension.

We then used the osmotic perturbation to constrain the membrane tension independently of the Piezo1 cluster measurements. The isotonic condition was assigned an effective membrane tension of approximately 0.2 pN/nm, whereas exposure to 120-mOsm Ringer’s solution was estimated to increase the tension to approximately 0.6 pN/nm (Fig. 4d). Supplementary Note 6 describes this estimate using published tether-force measurements^45^ under hypo-osmotic stimulation and the relation between tether force and effective membrane tension. These tension values, together with the experimentally measured relative Piezo1 density changes, were then used directly as inputs to the Brownian-dynamics simulations without further adjustment to match cluster size. Increasing the tension from 0.2 to 0.6 pN/nm reshaped the pair potential and shifted the predicted Piezo1 organization toward larger finite assemblies. The resulting cluster-area distributions reproduced the experimentally observed increase after hypo-osmotic swelling (Fig. 4d,e), providing a predictive test of the tension-dependent clustering mechanism.

Together, these results show that hypo-osmotic swelling shifts Piezo1 organization through the same biophysical mechanism derived above. Increasing membrane tension shifts Piezo1 toward a flatter conformation, reduces the residual boundary mismatch, and reshapes the balance between short-range attraction and longer-range repulsion in the effective pair potential. This change lowers the density threshold for finite-cluster formation, so a Piezo1 population at experimentally measured density is driven into the clustered regime. The agreement between the observed MINFLUX cluster-area distributions and the tension-shifted Brownian-dynamics simulations therefore provides a perturbative test of the model: changing membrane tension alone is sufficient to predict the direction and magnitude of Piezo1 cluster growth after swelling.

**Figure 4:**
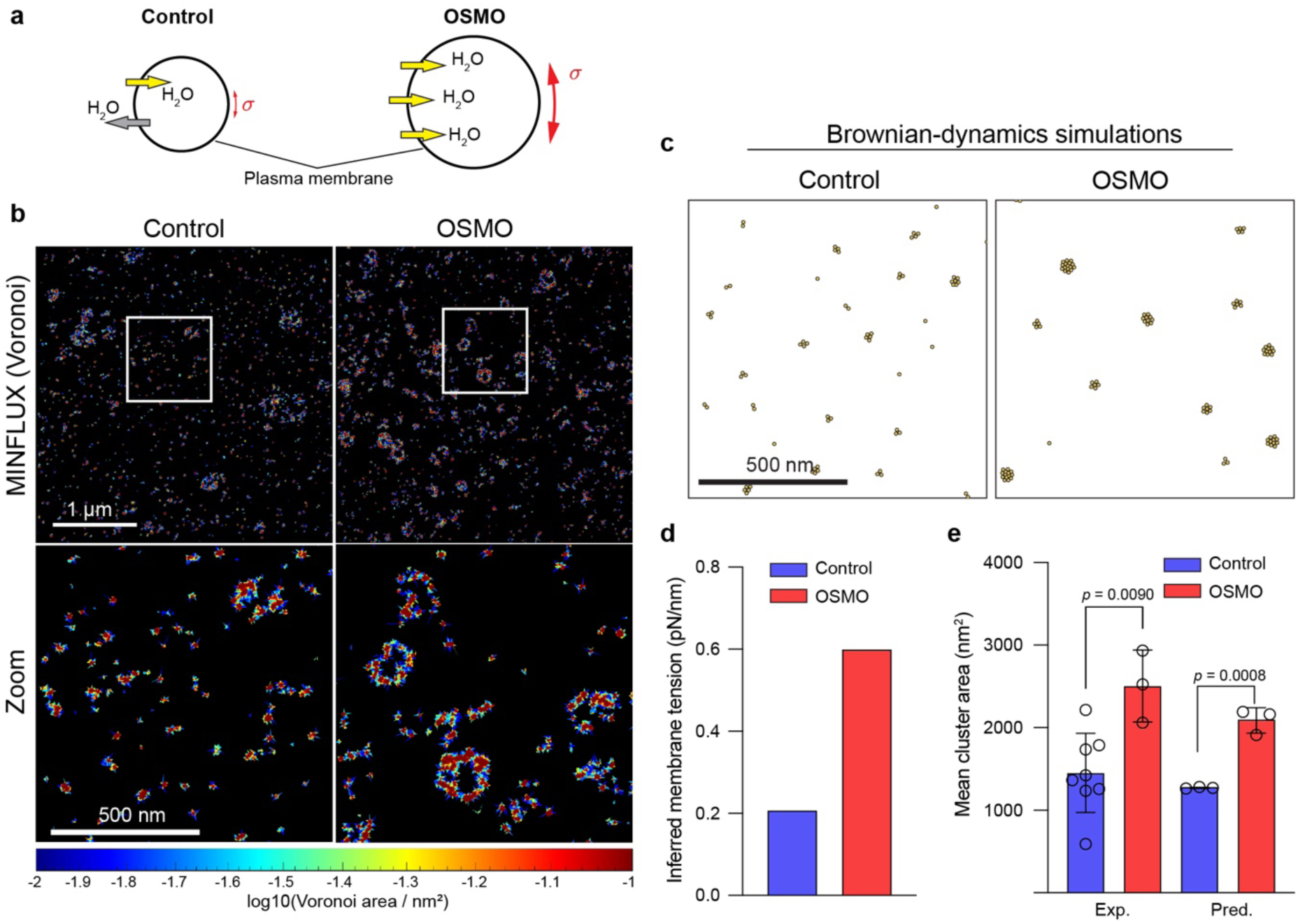
Hypo-osmotic swelling provides a perturbative test of tension-driven Piezo1 cluster growth. **a**, Schematic of control and hypo-osmotic conditions (OSMO). Water influx under hypo-osmotic treatment increases membrane tension and is predicted to shift Piezo1 organization toward larger clusters. **b,** MINFLUX imaging of Piezo1 under control and hypo-osmotic conditions, displayed as Voronoi-density maps with magnified views of representative regions. Hypo-osmotic treatment increases the size and compactness of Piezo1 assemblies. **c,** Model predictions for Piezo1 organization under control and hypo-osmotic conditions with channel-side parameters held fixed, shown at the same qualitative scale as the experimental comparison. **d,** Effective membrane tension inferred from the applied osmotic conditions, increasing from approximately 0.2 pN/nm under isotonic conditions to approximately 0.6 pN/nm following exposure to 120-mOsm Ringer’s solution. These values provide the mechanical inputs for the simulations in c and e. **e,** Experimental (Exp.) and predicted (Pred.) mean Piezo1 cluster area under control and hypo-osmotic conditions. Simulations performed using the membrane tensions in d reproduce the experimentally observed increase in cluster size without adjusting tension to the cluster measurements (Exp. Control n = 8, OSMO n = 3; Pred. n = 3; statistical significance was determined by two-tailed Student’s t-test). Together, the agreement between predicted and measured cluster growth provides a forward test of the tension-dependent clustering mechanism.

### Piezo1 density and cluster geometry map cell types onto a common membrane organization landscape

Having established that Piezo1 density and membrane tension regulate clustering within individual systems, we next asked whether measurements from distinct cellular systems could be placed on the same model-derived organization landscape. To do so, we compiled published estimates of Piezo1 abundance and surface density from complementary measurement modalities, including immunofluorescence imaging, patch-clamp-based density estimates, and combined imaging/electrophysiology datasets^17,18,22,46–48^ (Fig. 5a,b). The Human Protein Atlas (HPA) measurements in Fig. 5a provide a qualitative context for differences in Piezo1 expression across cell types but are not used as quantitative inputs to the model. These measurements span a broad experimental range, from sparse Piezo1 distributions in red blood cells to higher-density regimes in Neuro-2a and HEK293T cells. For systems with both density estimates and nanoscale organization measurements, we then compared the measured cluster geometry against the model predictions (Fig. 5c-e). Density fixes the channel-abundance axis of the model, whereas measured cluster size provides an independent structural constraint on how far each system has progressed from dispersed channels toward finite assemblies (Fig. 5f). This framework therefore tests whether Piezo1 organization states observed across different cellular systems can be explained by the same density-tension landscape rather than by channel abundance alone.

**Figure 5:**
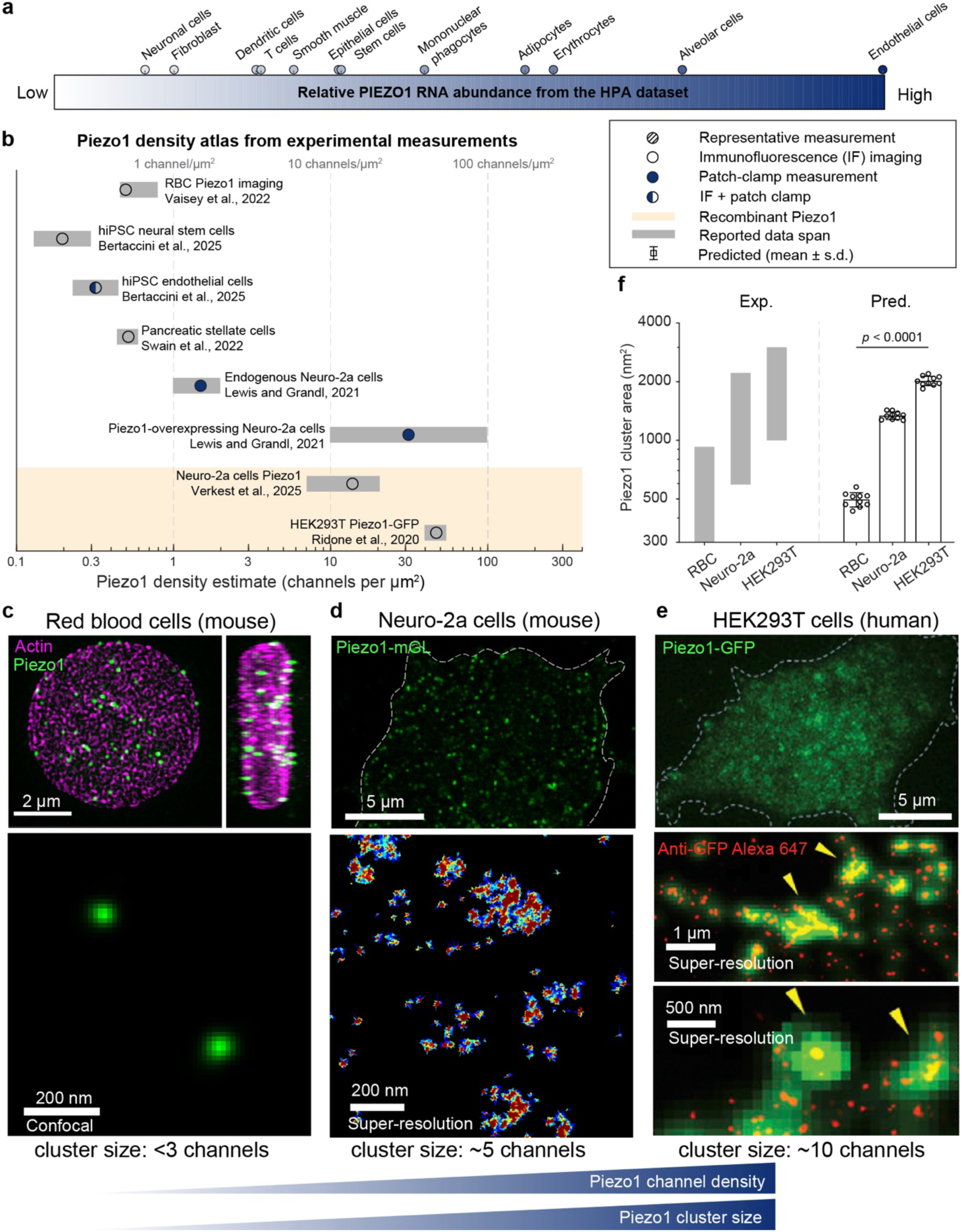
A Piezo1 density atlas positions distinct cell types on a common organization landscape. **a**, Relative Piezo1 abundance across representative cell types from the Human Protein Atlas, shown as a qualitative expression landscape spanning blood, immune, stromal, epithelial, and endothelial lineages. These measurements are not used as quantitative inputs to the model. **b,** Atlas of published Piezo1 density estimates compiled from immunofluorescence imaging, patch-clamp measurements, or combined datasets. Sources are Ref. [22] for red blood cells, Ref. [47] for hiPSC-derived neural stem cells and endothelial cells, Ref. [48] for pancreatic stellate cells, Ref. [17, 46] for endogenous and overexpressed Neuro-2a cells, and Ref. [18] for stable HEK293T Piezo1-GFP cells. These measurements define the experimentally relevant density range for comparing red blood cells, Neuro-2a cells, and HEK293T cells. **c-e,** Representative imaging phenotypes for red blood cells, Neuro-2a cells, and HEK293T cells. Sparse Piezo1 channels in red blood cells are consistent with the dispersed regime, whereas larger Piezo1 clusters in Neuro-2a and HEK293T cells are consistent with finite clustering at progressively higher density. Insets show higher-magnification or super-resolution views used to estimate cluster size. **f,** Experimentally measured (Exp.; left) and predicted (Pred.; right) Piezo1 cluster area across the representative systems, showing the progression from small assemblies in red blood cells to intermediate finite clusters in Neuro-2a cells and larger finite assemblies in HEK293T cells. (Pred. n = 10; statistical significance was determined by one-way ANOVA followed by Tukey’s multiple comparisons test)

Across the compiled datasets, the measurements reveal a graded, but not strictly linear, progression in Piezo1 organization (Fig. 5b,f). Red blood cells occupy the low-density end of the comparison, with published estimates near 0.5-1 Piezo1 channels per μm^2^ and apparent assemblies containing fewer than three channels^22^. Their measured cluster areas remain below ∼500 nm^2^, consistent with a sparse distribution close to the isolated-channel or small-oligomer limit. Neuro-2a cells^46^ span an intermediate range, with endogenous estimates approximately 1-3 channels per μm^2^ and overexpression datasets extending toward approximately 30 μm^-2^. Nanoscale imaging reveals representative finite assemblies containing approximately five channels with characteristic areas near 1,300 nm^2^. HEK293T Piezo1-GFP cells^18^ occupy a higher-density clustered regime, with reported densities near 10 channels per μm^2^ and characteristic assemblies of approximately 10 channels with areas near 2,000 nm^2^. Thus, despite differences in labeling strategy and measurement modality, the representative systems follow the qualitative sequence predicted by the model: sparse dispersed channels give way to intermediate finite clusters and then to larger, but still finite, Piezo1 assemblies.

We then used the combination of measured density and cluster geometry to place each system on the model-derived membrane-organization landscape (Fig. 5f). Because the same channel density can produce different cluster sizes at different membrane tensions, density alone does not uniquely determine organization state. For each cell type, the experimentally reported Piezo1 density was therefore fixed in Brownian-dynamics simulations, membrane tension was varied, and the predicted cluster geometry was compared with the corresponding experimental measurement. The range of tensions that reproduced the observed organization was reported as the compatible membrane-context range rather than as a direct mechanical measurement. Red blood cells admit a broad compatible range because their small cluster sizes remain close to the dispersed limit and therefore provide only a weak geometric constraint. Neuro-2a and HEK293T cells map to narrower sub-opening tension ranges of approximately 0.5-0.7 pN/nm and 0.3-0.5 pN/nm, respectively. These values lie below the ∼1-2 pN/nm single-channel opening scale used to parameterize the coarse-grained channel model, indicating that collective Piezo1 organization can emerge before full isolated-channel activation. The cross-system comparison therefore provides a consistency test of the density-tension framework, as heterogeneous cellular measurements can be positioned on the same organization landscape using density as the experimental input and cluster geometry as the independent structural constraint.

### Receptor-associated membrane remodeling recruits Piezo1 and nucleates finite clusters

The density-tension framework developed above treats the membrane as a spatially uniform mechanical background. Living plasma membranes, however, contain substantial local heterogeneity. Receptor assemblies and lipid-order domains reorganize membrane composition and protein partitioning^18,49^, while cortical attachment, force-bearing adhesions^19^, membrane curvature, and protein crowding create additional local mechanical cues that reshape the environment sensed by Piezo1^20,30^. We therefore asked whether local membrane remodeling can recruit Piezo1 strongly enough to drive an otherwise subcritical population into the finite-cluster regime derived above. Within this formulation, membrane heterogeneity determines where Piezo1 accumulates, whereas the previously derived Piezo1-Piezo1 interaction determines whether the accumulated channels form a stable finite cluster (Fig. 6a).

We denote the in-plane position of a Piezo1 channel by ***R*** and the center of a mechanically remodeled receptor domain by ***x****_R_*. We represent the local mechanical perturbation as

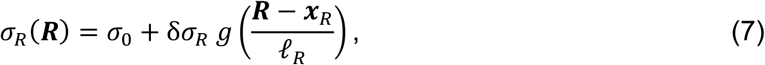

where *σ*_0_ is the spatially uniform membrane tension, while δ*σ_R_* and *P_R_* set the amplitude and spatial range of the perturbation. Importantly, δ*σ_R_* does not represent a literal local increase in membrane tension, but rather a tension-equivalent measure of the additional mechanical bias experienced by Piezo1, corresponding to the membrane tension that would produce the same change in Piezo1 free energy as the local membrane remodeling. This additive representation is motivated by continuum membrane theories in which domain-specific mechanical constraints can introduce tension-like contributions to the membrane free energy^50,51^, with δ*σ_R_* playing the analogous role while the physical membrane tension *σ*_0_ remains spatially uniform. The dimensionless function *g* describes the spatial profile of the remodeled domain, with *g*(0) = 1 at the domain center and *g*(∞) = 0 far from the domain. The field *σ_R_*(***R***) therefore approaches *σ*_0_ + δ*σ_R_* at the center and returns to the uniform background value *σ*_0_ outside the remodeled region. Written in this form, the field coarse-grains local effects of lipid organization, curvature, membrane-cortex coupling, receptor-associated crowding, and related membrane remodeling into a common mechanical bias that couples to the Piezo1 projected area.

We couple this local field to the same single-channel mechanics developed above,

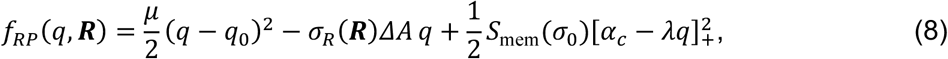

where *q* is the membrane-coupled Piezo1 conformational coordinate and Δ*A* is the projected-area gain associated with flattening. The local field *σ_R_*(***R***) changes the energetic balance between Piezo1 conformation, projected-area expansion, and the residual boundary mismatch at each membrane position. This construction does not introduce new receptor-Piezo1 binding energy, so local membrane remodeling changes the energetic cost of placing the same mechanically adaptive Piezo1 channel at different membrane positions.

We next minimize Eq. (8) over *q* to obtain the effective one-body recruitment potential

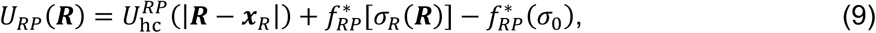

where 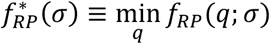. A region with *U_RP_* < 0 therefore preferentially recruits Piezo1. For weak perturbations, 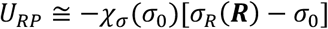, where 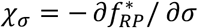 defines the mechanical susceptibility of an isolated channel. Supplementary Note 7 derives this weak-field response by expanding the minimized one-body free energy about the far-field state, showing that the same mechanical susceptibility that governs Piezo1 conformational response also determines its recruitment by a localized membrane perturbation. We include an additional hard-core contribution when the remodeled structure is sterically inaccessible to Piezo1.

The recruitment potential maps this mechanical heterogeneity onto the local Piezo1 density. In the dilute pre-nucleation regime, 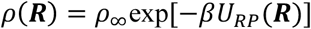, where *ρ*_∞_ is the far-field channel density (Fig. 6b,c). Because we keep the Piezo1-Piezo1 pair potential fixed at the far-field membrane state, the relevant clustering-onset density, *ρ*_on_(*σ*_0_), is the same onset obtained above for a uniform membrane at tension *σ*_0_ for the fixed channel parameters used here. An effective mechanically remodeled domain therefore nucleates a finite cluster when recruitment raises the local density from *ρ*_∞_ to *ρ*_on_ . Expressed as a free-energy threshold, the required critical recruitment gain is

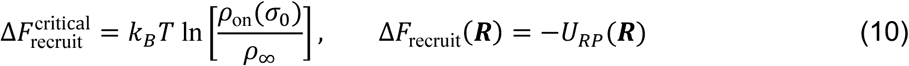

The model predicts local clustering where 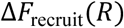 exceeds this threshold. Equation (10) captures the competition between the local recruitment free-energy gain supplied by membrane remodeling and the logarithmic density deficit ln[*ρ*_on_/*ρ*_∞_], which measures how far the far-field Piezo1 population lies below the pre-existing clustering boundary. A remodeled membrane domain can therefore drive local clustering by supplying enough recruitment free energy to overcome this density deficit and move the local population across the clustering boundary established by the uniform-membrane theory.

We tested this mechanism using Brownian-dynamics simulations while holding the Piezo1-Piezo1 pair potential fixed at the far-field membrane state (Fig. 6d). The effective mechanical recruitment field acted only through the one-body recruitment potential *U_RP_*. A weak field recruited Piezo1 and produced only diffuse enrichment because the recruitment gain remained below the threshold in Eq. (10). Increasing the field amplitude or spatial range increased the local Piezo1 density until the recruitment gain exceeded this threshold, allowing the pre-existing short-range attraction and longer-range repulsion to stabilize a finite assembly. The simulations show how a spatially localized recruitment field converts a subcritical Piezo1 population into a finite cluster once the local density crosses the uniform-membrane onset without introducing an additional receptor-mediated attraction between channels.

We next compared this prediction with fluorescence images published by Geng et al.^52^, in which Piezo1 and Toll-like receptor 4 (TLR4) were visualized in bone-marrow-derived macrophages under control conditions and following stimulation with lipopolysaccharide (LPS). We quantified Piezo1 enrichment using the receptor-centered radial enrichment analysis described in Methods. Following LPS stimulation, Piezo1 was more strongly enriched around TLR4-associated regions and the radial intensity profile broadened (Fig. 6e-g). Increasing the amplitude or spatial range of the effective recruitment field produced the same qualitative redistribution in the model. Because *σ_R_*(***R***) coarse-grains membrane-level changes in lipid organization, adaptor recruitment, protein crowding, and cortex coupling, we treat it as an effective mechanical field rather than assign it to a single microscopic source. Within this description, receptor-associated remodeling lowers the local free energy of Piezo1, promotes channel accumulation, and raises the local density toward the pre-existing clustering boundary. Once this boundary is crossed, the same Piezo1-Piezo1 interaction derived for the uniform membrane stabilizes a finite receptor-associated assembly.

These results extend the density-tension framework from uniform to spatially patterned membranes. In a uniform membrane, mechanics determines whether a Piezo1 population enters the finite-cluster regime, while in a heterogeneous membrane, local mechanics additionally determines where that transition occurs. Membrane remodeling can therefore position finite Piezo1 assemblies at receptor-associated regions without increasing global channel abundance, providing a physical mechanism for spatially localizing mechanosensitive signaling.

**Figure 6:**
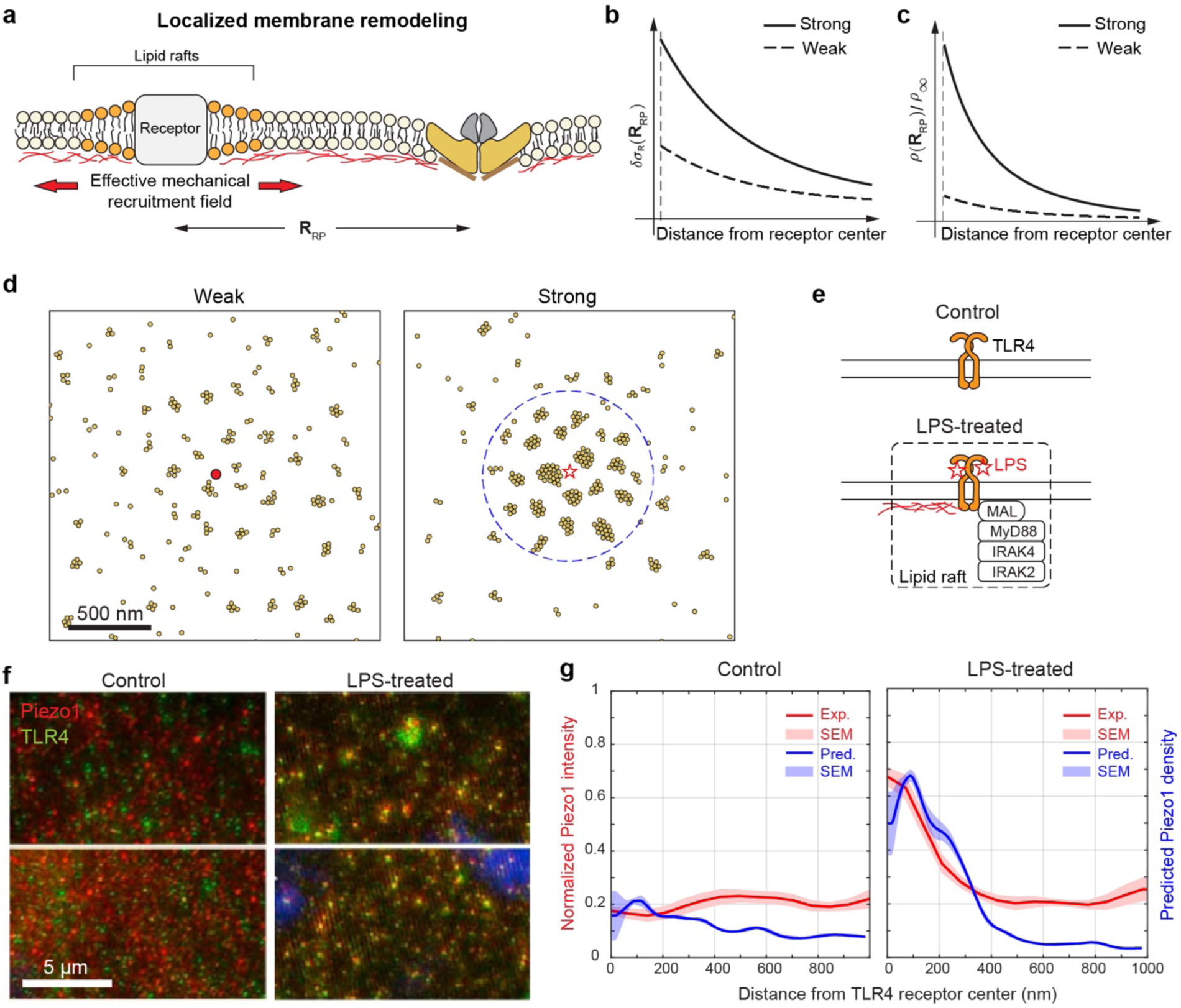
Receptor-associated local membrane remodeling recruits Piezo1 and promotes receptor-associated finite clustering. **a**, Schematic of an effective mechanically remodeled membrane domain generated by receptor organization, lipid remodeling, and cortex coupling. Their combined effect is represented by an effective local mechanical field *σ_R_*(***R***) superimposed on the far-field tension *σ*_O_. This field changes the minimized Piezo1 free energy and biases channels toward the remodeled region. **b,** Radial profiles of weak and strong effective mechanical recruitment fields as functions of distance from the receptor center. Increasing the field amplitude and spatial range extends the membrane region over which Piezo1 recruitment is energetically favored. **c,** Predicted radial Piezo1 enrichment around weak and strong recruitment fields. Stronger recruitment increases the local Piezo1 density while far-field density remains unchanged. **d,** Brownian-dynamics simulations illustrating the transition from diffuse recruitment to finite clustering. The localized field acts only as a one-body positional bias, while Piezo1 channels interact through the same pair potential evaluated at the far-field membrane state. A weak field (solid circle) produces local enrichment below the clustering threshold, whereas a stronger field (hollow star) raises the local density sufficiently for the existing Piezo1-Piezo1 interaction to stabilize finite assemblies. **e,** Schematic of TLR4 activation by lipopolysaccharide (LPS), illustrating receptor-associated changes in lipid organization, adaptor recruitment, and cortex coupling that can remodel the local membrane environment. **f,** Representative fluorescence images of Piezo1 and TLR4 under control and LPS-treated conditions^52^. **g,** Radial Piezo1 intensity profiles around TLR4-associated regions under control and LPS-treated conditions. Experimental measurements are compared with model profiles, showing that enhanced Piezo1 enrichment after LPS stimulation is consistent with a stronger or longer-ranged local recruitment field. (Exp. n = 10; Pred. n = 3)

## Discussion

This study establishes recursive membrane feedback as a physical mechanism by which Piezo1 converts force-dependent changes in channel shape into collective membrane organization. The central advance is that Piezo1 is not treated as a passive membrane inclusion with a prescribed geometry, but as a mechanically adaptive component whose conformation and surrounding membrane-cortex deformation are mutually determined. Membrane tension and membrane-cortex mechanics bias Piezo1 toward a flatter, more open-like state, reducing the deformation imposed on the surrounding membrane and changing how strongly it mechanically influences nearby Piezo1 proteins. Nearby channels then respond to this altered membrane shape, creating a feedback loop in which the conformation of one Piezo1 molecule affects the conformation and mechanical influence of its neighbors.

When neighboring channels share the same membrane deformation field, recursive feedback generates two competing interactions. Overlap of residual membrane deformations carries an elastic cost and produces longer-range repulsion, whereas at shorter separations, neighbor-induced flattening reduces the boundary mismatch and lowers the pair energy. This local relaxation is consistent with molecular simulations of crowded Piezo1 channels, in which overlapping membrane footprints promote flattening of the Piezo1 arms and pore widening^30^. The resulting short-range attraction and longer-range repulsion provide the physical basis for finite clustering, with attraction stabilizing nearby channels and the outer repulsive barrier limiting continued growth. The stability of this regime depends on conformational feedback strength and membrane tension, which shape the attractive basin and clustering threshold (Supplementary Figs. 2-4), while channel density determines how frequently diffusing Piezo1 molecules encounter and remain within that basin. Together, these effects link single-channel mechanosensitivity to the onset of collective Piezo1 organization.

Many-body Brownian-dynamics simulations confirm that these pair-level interactions are sufficient to generate finite Piezo1 organization. Increasing channel density favors finite assemblies, while higher membrane tension lowers the density threshold for clustering by reshaping the pair potential. The simulations therefore define a density-tension organization landscape in which Piezo1 can occupy dispersed or finite-cluster states depending on the combined channel supply and mechanical context of the membrane. Comparison with Piezo1-GFP organization in HEK293T cells provides an experimental test of this regime^18^. Using the measured Piezo1 channel density to constrain the simulations, the model reproduces not only the characteristic cluster size but also the broad, right-skewed distributions of cluster area and perimeter. Agreement across these geometric measures supports the interpretation that Piezo1 clusters represent a mechanically selected finite-cluster state rather than nonspecific membrane accumulation.

Hypo-osmotic swelling provides a complementary perturbative test of membrane tension. Swelling increases membrane tension while leaving the intrinsic channel-side parameters almost unchanged on the experimental timescale, allowing the model to predict how Piezo1 organization should shift under an independently imposed mechanical perturbation. Analysis of published super-resolution MINFLUX data from Neuro-2a cells shows larger Piezo1 assemblies under hypo-osmotic conditions, and Brownian simulations using tension-shifted pair potentials reproduce the increase in mean cluster area, supporting membrane tension as a causal regulator rather than simply a fitted correlate. More broadly, published density and imaging measurements place red blood cells near the sparse, dispersed limit, Neuro-2a cells in an intermediate finite-cluster regime, and HEK293T cells in a higher-density regime with larger but still finite assemblies. Despite differences in measurement modality, this progression is consistent with the density-tension framework and supports a common physical picture in which channel abundance and membrane mechanics jointly shape Piezo1 organization across cellular systems.

A key implication of our analysis is that Piezo1 collective organization can emerge at membrane tensions below the scale typically associated with full isolated-channel activation. Recent structural work provides complementary evidence that Piezo1 conformational remodeling can precede full pore opening, with membrane forces driving the channel toward a flatter closed state before full conductive activation^38,53,54^. In the model, sub-opening membrane tensions can still reshape the residual boundary mismatch and alter the attraction-repulsion balance between channels. Thus, Piezo1 clustering may precede, bias, or spatially organize channel activation rather than merely following from it. This provides a mechanism by which cells could tune mechanotransduction sensitivity without globally opening all channels. By changing membrane tension, cortex coupling, lipid organization, or local channel density, cells may reposition Piezo1 within the organization landscape and thereby regulate where mechanical signals are most likely to be converted into Ca^2+^ entry.

Local membrane heterogeneity further extends this mechanism from uniform self-organization to spatially patterned recruitment. Receptor-associated domains, lipid-ordered regions, cortical attachment sites, protein-crowded patches, and force-bearing adhesions can alter local membrane mechanics and create energetic sinks for Piezo1. Once recruitment raises the local channel density above the same clustering threshold that governs the uniform membrane, recursive Piezo1-Piezo1 interactions stabilize finite assemblies. The LPS-activated TLR4 analysis supports this sequence, with receptor-associated membrane remodeling increasing local Piezo1 enrichment and the radial enrichment profiles consistent with stronger local mechanical fields after stimulation, suggesting that local membrane mechanics can determine where finite clusters form without altering global channel abundance.

The spatial recruitment and clustering of Piezo1 may also help match Ca^2+^ entry to local signaling demand. Because intracellular Ca^2+^ is strongly buffered, the high Ca^2+^ concentration generated near an open channel falls rapidly with distance, making proximity between Ca^2+^ sources and downstream effectors important^55^. Clustering Piezo1 can therefore concentrate Ca^2+^ influx at sites where it is needed, consistent with discrete Ca^2+^ signals from Piezo1 assemblies and force-dependent recruitment of Piezo1 to focal adhesions^17,19^. In our framework, such positioning is produced by a localized mechanical recruitment field, whose amplitude δ*σ_R_* and range *P_R_* specify the strength and spatial extent of membrane remodeling. Force-bearing adhesions and receptor-associated domains provide experimentally established examples of structures that locally reorganize membrane mechanics and recruit Piezo1^19,49,52^. Mitochondria can likewise position near sites of Ca^2+^ entry to shape local Ca^2+^ signaling^56,57^, while ER-plasma-membrane contacts provide highly localized membrane domains specialized for Ca^2+^ exchange^58^. These observations suggest a broader spatial organization principle in which mechanically patterned membrane regions position Piezo1 close to sites of Ca^2+^ utilization or handling. What remains unknown is whether Piezo1 clustering is required to generate sufficient local Ca^2+^ signals at these sites, and whether organelle-associated membrane remodeling can in turn recruit or stabilize Piezo1 clusters. These possibilities could be tested by correlating Piezo1 organization and organelle position with local Ca^2+^ signals in the same cells.

This framework differs from previous continuum and membrane-inclusion models by treating Piezo1 shape as a dynamic mechanical coordinate rather than a fixed boundary condition^41^. Wiggins and Phillips developed an analytic framework for understanding MscL gating based on lipid-protein interaction energies, showing how bilayer mechanics contributes to free-energy differences between channel states^31,37^. Ursell et al. extended this picture to cooperative gating and attraction between MscL-like channels through bilayer-thickness deformations around fixed inclusions^32^. Because these deformations arise largely from hydrophobic mismatch, the resulting interactions are short-ranged and decay over molecular length scales set by the bilayer thickness. Conical-inclusion theories, including the work of Weikl^41^, describe a different fixed-geometry limit in which curvature-mediated interactions are controlled by membrane tension. Piezo-specific membrane-footprint models developed by Haselwandter and colleagues^25,28,29^ brought this continuum perspective to Piezo channels by showing how a prescribed curved Piezo dome deforms the surrounding membrane and contributes to tension sensitivity. Across these approaches, however, channel geometry is specified in advance through a fixed thickness mismatch, cone angle, contact curvature, or Piezo footprint. By allowing Piezo1 shape itself to respond to membrane mechanics, the present framework makes the membrane footprint state-dependent, so that channel conformation, membrane deformation, and interactions between neighboring channels are determined together.

This coarse-grained description focuses on the core membrane-mediated mechanism and therefore leaves additional molecular and cellular complexity unresolved. Force-dependent flattening is represented by a single coordinate that captures how channel geometry reshapes the surrounding membrane, rather than the full kinetics of pore opening, inactivation, and Ca^2+^ conduction. The Brownian-dynamics simulations also approximate many-channel interactions as pairwise additive. In crowded membranes, simultaneous overlap of multiple deformation fields may produce non-additive conformational and membrane responses that are not captured by the two-channel potential, potentially shifting the quantitative stability and geometry of finite clusters. Membrane tension is also inferred in several comparisons from channel density and cluster geometry rather than measured directly, and the published datasets differ in labeling strategy, imaging modality, and expression level. Sensitivity analyses of the channel geometry and mechanical parameters show that these parameter variations shift the phase boundaries quantitatively while preserving the finite-cluster regime (Supplementary Figs. 8 and 9). These limitations therefore shift where individual systems fall within the organization landscape without altering the existence of the finite-cluster regime.

The present framework considers an idealized, locally flat membrane, whereas cellular membranes exhibit complex geometries and spatially varying mechanical environments. Extending the model to cell-scale membrane curvature could reveal how local curvature and tension jointly influence Piezo1 conformation and collective recruitment. This may be particularly relevant to processes such as keratinocyte migration and wound healing, where Piezo1 becomes spatially enriched within migrating cells and wound edges^59^. Beyond this extension, the density-tension landscape suggests direct experimental tests. Perturbing Piezo1 conformational responsiveness, membrane tension, membrane-cortex stiffness, or lipid and receptor organization should shift clustering or local enrichment. Simultaneous measurements of membrane mechanics, Piezo1 nanoscale organization, and Ca^2+^ influx would further test whether mechanically selected clusters correspond to functional signaling hotspots.

Together, by making the Piezo1 membrane footprint both tension-dependent and collective, the framework creates recursive feedback in which membrane tension changes channel shape, channel shape changes the surrounding membrane deformation field, and that field reshapes the interactions experienced by neighboring channels. The membrane footprint therefore becomes a state-dependent interaction field linking channel flattening to non-monotonic pair interactions, finite clustering, and many-body spatial organization. This self-consistent coupling connects isolated-channel energetics to finite mesoscale clusters and local recruitment at mechanically patterned membrane domains, explaining how Piezo1 can remain dispersed, assemble into finite clusters, or become enriched at receptor- or adhesion-associated structures depending on membrane context. More broadly, the organization of mechanosensitive channels may represent a mechanically tunable variable that couples membrane remodeling to localized mechanotransduction across diverse cellular environments.

## Methods

### Brownian-dynamics simulations

Many-body organization was simulated using overdamped Brownian-dynamics in two dimensions with periodic boundary conditions. For channel *i* at position **R***_i_*, the equation of motion is

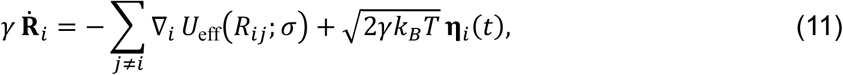

where *γ* is the translational drag coefficient, 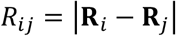, and **η***_i_*(*t*) is Gaussian white noise with zero mean and unit variance. The corresponding single-particle diffusivity is *D* = *k_B_T*/*γ*. Simulations were implemented in custom MATLAB scripts.

For a simulation box of side length *L* containing *N* channels, the mean areal density is 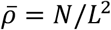. Density sweeps were performed by varying *N* and/or *L* at fixed membrane tension, whereas tension sweeps were performed by changing *σ* in the pair potential while holding the channel-side parameters fixed. Random initial conditions were generated with hard-core overlap exclusion, and simulations were advanced until cluster statistics reached a stationary distribution. Independent realizations were used to estimate the mean cluster size, cluster fraction, and cluster diffusivity under each condition. To quantify the onset of clustering in simulation, we monitored the fraction of channels belonging to clusters of size three or larger. Clusters were identified from the particle coordinates using a parameter-locked spatial clustering rule applied consistently within each sweep. The simulated onset density was defined operationally as the density at which this clustered fraction crossed the threshold shown in Supplementary Fig. 5. For the finite-cluster regime, the same trajectories were used to calculate the mean number of channels per cluster and the dependence of effective cluster diffusivity on cluster occupancy.

For direct comparison with confocal imaging, Brownian-dynamics particle configurations were converted to confocal-like intensity maps by convolving the particle positions with an isotropic point-spread kernel at the experimental imaging scale. The resulting synthetic images were then segmented using the same geometric pipeline used for the experimental HEK293T data, so that cluster area and perimeter were extracted consistently from simulations and experiments.

### Model-constrained inference of membrane context from Piezo1 organization

The comparison between theory and experiment followed the workflow summarized in Supplementary Fig. 6. Channel-side parameters were fixed from structural and biophysical measurements before comparison with cellular organization data. Piezo1 density and membrane tension were then treated as physical control variables. For perturbative comparisons, such as hypo-osmotic swelling, the model tested whether changing the membrane mechanical state alone shifted the cluster-size distribution in the predicted direction while keeping the channel-side parameters fixed. To assess the robustness of this inference procedure, we varied the geometric and mechanical parameters across the ranges shown in Supplementary Figs. 8 and 9, which quantify how these variations shift the organization boundaries.

For cross-system comparisons, the experimentally measured Piezo1 density *ρ*_exp_ was used as an input constraint. At a trial membrane tension *σ*, Brownian-dynamics simulations were performed at the observed density to generate a predicted cluster distribution. Let *A*_sim_∼*σ*; *ρ*_exp_p denote the simulated cluster area and *A*_exp_ the measured cluster area extracted from the corresponding imaging data. The membrane context was estimated by minimizing the mismatch:

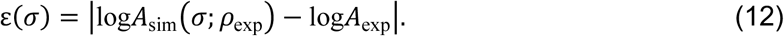

Equivalently, membrane tension was updated iteratively until the simulated and measured cluster scales agreed within the chosen tolerance. In this way, red blood cells, Neuro-2a cells, and HEK293T cells were placed on the same theoretical organization landscape using experimentally constrained density and cluster size, while the inferred membrane tensions were interpreted as model-derived membrane-context ranges rather than as direct mechanical measurements. The sensitivity analyses in Supplementary Figs. 8 and 9 show that, although the precise inferred ranges shift quantitatively with the assumed parameter set, the ordering of systems on the organization landscape and the existence of a finite-cluster regime remain robust.

### Receptor-centered radial enrichment analysis

Fluorescence images of Piezo1 and TLR4 were analyzed using custom MATLAB scripts. TLR4-positive receptor-associated regions were identified from the TLR4 fluorescence channel and used as spatial reference positions for the Piezo1 analysis. For each receptor-associated region, Piezo1 fluorescence intensity was measured as a function of radial distance from the TLR4-defined center to generate a receptor-centered radial intensity profile. Piezo1 intensity was normalized to the far-field signal, allowing enrichment near TLR4-associated regions to be compared across images and experimental conditions. Radial profiles were averaged across analyzed receptor-associated regions separately for control and LPS-treated cells and were compared with the radial enrichment profiles predicted by the localized membrane-field model. The same MATLAB analysis parameters were applied to control and LPS-treated images.

## Data and code availability

The experimental imaging data analyzed in this study were obtained from previously published datasets cited in the manuscript. Processed data, simulation outputs, and the MATLAB code used for Brownian-dynamics simulations are available at our GitHub repository: https://github.com/zxguo98/Piezo1_Membrane_Organization.

## Acknowledgements

This work was supported by NIH Award U54CA261694 (V.B.S.); NSF CEMB Grant CMMI-154857 (V.B.S.); NSF Grant DMS-2347834 (V.B.S.); National Institute of Biomedical Imaging and Bioengineering (NIBIB) Awards R01EB017753 (V.B.S.) and R01EB030876 (V.B.S.); National Institute of General Medical Sciences award R01GM155943 (V.B.S.) and National Institute of Diabetes and Digestive and Kidney Diseases grant R01DK144619 (V.B.S.). The authors thank Beatriz Hernaez Estrada for her insightful discussions and feedback during the early development of the study, particularly in exploring potential mechanisms of Piezo1 function in immune cells. The authors gratefully acknowledge the valuable comments and suggestions from Vivek Sharma and Vinayak.

## Author contributions

V.B.S. conceived the project and derived the theoretical framework. Z.G. developed the numerical analysis and analyzed the experimental data. Z.G. and A.B. conducted the quantitative analysis of super-resolution and confocal images. V.B.S. led the drafting of the manuscript. Z.G. contributed substantially to manuscript preparation, including preparation of figures, and A.B., M.Dh., and M.De. contributed to revisions and proofreading.

