## Supplementary Information for "Recursive feedback between Piezo1 conformation and membrane mechanics drives self-organization into finite clusters"

for

### Supplementary Note 1: Elimination of the membrane field for a single channel

For one channel centered at the origin on a flat background, the starting free-energy functional is

$$F_1[u,q;\sigma] = \int_{r>a_{\mathrm{ch}}} \mathrm{d}^2 r \left[\frac{\kappa_{\mathrm{eff}}}{2}(\nabla^2 u)^2 + \frac{\sigma}{2}|\nabla u|^2\right] + \frac{\mu}{2}(q-q_0)^2 - \sigma \Delta A\, q, \tag{S1}$$

where $u(\mathbf{r})$ is the channel-induced membrane deformation, $q$ is the channel flattening coordinate, $\kappa_{\mathrm{eff}}$ is the effective membrane-cortex bending stiffness, $\sigma$ is the membrane tension, $\mu$ is the bare stiffness of the internal coordinate, $q_0$ is the reference state, and $\Delta A$ is the projected area gain associated with flattening. The channel imposes the state-dependent boundary mismatch $\partial_r u(a_{\mathrm{ch}}) = \alpha(q) = [\alpha_c - \lambda q]_+$, with effective channel radius $a_{\mathrm{ch}}$, closed-state mismatch $\alpha_c$, and feedback strength $\lambda$. Variation with respect to $u$ gives the screened bending equation

$$\kappa_{\mathrm{eff}}\nabla^4 u - \sigma \nabla^2 u = 0, \qquad \xi = \sqrt{\frac{\sigma}{\kappa_{\mathrm{eff}}}}, \tag{S2}$$

together with the decaying axisymmetric solution $u(r) = A\, K_0(\xi r)$, where $K_0$ is the modified Bessel function of the second kind. Using $\partial_r K_0(\xi r) = -\xi K_1(\xi r)$ and imposing the boundary condition at $r = a_{\mathrm{ch}}$ gives

$$A = -\frac{\alpha(q)}{\xi K_1(z)}, \qquad z = \xi a_{\mathrm{ch}}, \tag{S3}$$

so that the minimizing deformation field is

$$u^*(r;q,\sigma) = -\frac{\alpha(q)}{\xi K_1(z)} K_0(\xi r). \tag{S4}$$

Substituting $u^*$ back into the membrane energy yields the on-shell elastic cost

$$F^*_{\mathrm{mem}}(q;\sigma) = \frac{1}{2} S_{\mathrm{mem}}\, \alpha(q)^2, \qquad S_{\mathrm{mem}} = 2\pi\kappa_{\mathrm{eff}} z\, \frac{K_0(z)}{K_1(z)}. \tag{S5}$$

The reduced one-body free energy is therefore

$$f_1(q;\sigma) = \frac{\mu}{2}(q-q_0)^2 - \sigma\Delta A\, q + \frac{1}{2} S_{\mathrm{mem}}([\alpha_c - \lambda q]_+)^2 \tag{S6}$$

In the active branch, $q < q_{\mathrm{sat}}$ with $q_{\mathrm{sat}} = \alpha_c/\lambda$, the clipped law is linear and the free energy becomes

$$f_1^<(q;\sigma) = \frac{\mu_{\mathrm{eff}}}{2} q^2 - (\mu q_0 + \lambda\alpha_c S_{\mathrm{mem}} + \sigma\Delta A) q + \mathrm{const}, \qquad \mu_{\mathrm{eff}} = \mu + \lambda^2 S_{\mathrm{mem}}. \tag{S7}$$

Minimization then gives the preferred isolated-channel state

$$q_\infty(\sigma) = \frac{\mu q_0 + \lambda\alpha_c S_{\mathrm{mem}} + \sigma\Delta A}{\mu + \lambda^2 S_{\mathrm{mem}}}, \tag{S8}$$

provided this minimum remains below $q_{\mathrm{sat}}$. The corresponding residual mismatch is $\alpha_\infty(\sigma) = \alpha_c - \lambda q_\infty(\sigma)$. If the unconstrained minimum reaches or exceeds $q_{\mathrm{sat}}$, the preferred boundary mismatch saturates at zero and the one-body response crosses into the saturated branch. The background-curvature extension used only in the Supplementary Information is obtained by replacing the boundary condition with $\alpha(q) - a_{\mathrm{ch}} H_0$, where $H_0$ is the local mean curvature of the slowly varying background membrane. This shifts the preferred state and residual mismatch but does not modify the membrane-elimination procedure itself.

**Supplementary Note 2. Elimination of the membrane field for a channel pair**

For two channels with centers separated by $R = |\mathbf{R}_1 - \mathbf{R}_2|$, the free energy at fixed internal states $q_1$ and $q_2$ is

$$F_{12}[u, q_1, q_2; \sigma] = U_{\mathrm{hc}}(R) + \sum_{i=1}^{2} \left[\frac{\mu}{2}(q_i - q_0)^2 - \sigma \Delta A\, q_i\right] + \int_{\Omega} \mathrm{d}^2 r \left[\frac{\kappa_{\mathrm{eff}}}{2}(\nabla^2 u)^2 + \frac{\sigma}{2}|\nabla u|^2\right], \qquad \text{(S9)}$$

where $\Omega$ is the membrane exterior to the two channel footprints and $U_{\mathrm{hc}}(R)$ is the hard-core exclusion. Each channel imposes the boundary condition

$$\partial_{n_i} u|_{C_i} = \alpha(q_i) = [\alpha_c - \lambda q_i]_+, \qquad i = 1,2. \qquad \text{(S10)}$$

Because the governing equation and boundary conditions are linear in the membrane field, elimination of $u$ yields an exact quadratic boundary form. In the isotropic coarse-grained description and to first order in the weak-overlap regime, the reduced two-channel free energy may be written as

$$F_2(R; q_1, q_2) = U_{\mathrm{hc}}(R) + \sum_{i=1}^{2} f_1\,(q_i; \sigma) + J(R)\,\alpha(q_1)\alpha(q_2) - \lambda G(R)\,[q_1 \alpha(q_2) + q_2 \alpha(q_1)], \qquad \text{(S11)}$$

where $J(R)$ is the direct-overlap kernel and $G(R)$ is the feedback kernel. The large-separation form of these kernels is $J(R) = J_0 K_0(\xi R), G(R) = G_0 \xi K_1(\xi R)$, with $J_0 > 0$, $G_0 > 0$, and $\xi = \sqrt{\sigma/\kappa_{\mathrm{eff}}}$. Here, the constant $J_0 = 2\pi\kappa_{\mathrm{eff}}/K_1^2(\xi a_{\mathrm{ch}})$ sets the direct elastic cost of field overlap, where $a_{\mathrm{ch}}$ is the effective Piezo1 channel footprint radius. The quantity $G_0 = a_{\mathrm{ch}} J_0$ then parameterizes the magnitude of the neighbor-induced shift in the flattening coordinate.

In the symmetric channel-pair problem, $q_1 = q_2 = q$, so

$$F_2(R; q) = U_{\mathrm{hc}}(R) + 2 f_1(q; \sigma) + J(R)\alpha(q)^2 - 2\lambda G(R)\, q\, \alpha(q). \qquad \text{(S12)}$$

Expanding about the isolated optimum, $q = q_\infty + \delta q$ and $\alpha(q) = \alpha_\infty - \lambda \delta q$, and retaining only terms of order $\mathrm{e}^{-\xi R}$ gives

$$F_2(R; q) = 2 f_1(q_\infty; \sigma) + U_{\mathrm{hc}}(R) + J(R)\alpha_\infty^2 - 2\lambda G(R)\, q_\infty \alpha_\infty + \mu_{\mathrm{eff}}(\delta q)^2 + \mathcal{O}\big(\mathrm{e}^{-2\xi R}\big). \qquad \text{(S13)}$$

Minimization with respect to $\delta q$ contributes only at order $\mathrm{e}^{-2\xi R}$, so the leading effective pair interaction becomes

$$U_{\mathrm{eff}}(R; \sigma) = U_{\mathrm{hc}}(R) + B(\sigma) K_0(\xi R) - C(\sigma)\, \xi K_1(\xi R), \qquad \text{(S14)}$$

with $B(\sigma) = J_0 \alpha_\infty^2, C(\sigma) = 2\lambda G_0 q_\infty \alpha_\infty$. The $K_0$ term is repulsive because it is the direct elastic cost of overlapping membrane deformation fields. The $\xi K_1$ term is attractive because one channel locally flattens its neighbor, thereby reducing the boundary mismatch and lowering the pair energy. In the saturated branch, $\alpha_\infty = 0$, so both amplitudes vanish at leading order. A useful control is the low-feedback limit. As $\lambda$ is reduced toward zero, the flattening response no longer relaxes the imposed boundary mismatch, so the attractive contribution disappears and the interaction becomes effectively repulsive outside hard contact. Supplementary Fig. 3 shows this directly at the phase-diagram level. The same figure establishes that the hump-shaped finite-cluster regime is created by recursive state–shape feedback rather than by direct overlap alone. Supplementary Figs. 8 and 9 then show that varying $\alpha_c$, $\kappa_{\text{eff}}$, $\mu$, $a_{\text{ch}}$, and $\Delta A$ shifts the phase boundaries quantitatively but preserves the same three qualitative regimes. The finite-cluster window is therefore robust to broad mechanical and geometric parameter variation.

**Supplementary Note 3. Onset density and the origin of finite cluster-size distributions**

Let $d^*$ denote the minimum of $U_{\text{eff}}(R;\sigma)$ and $d_b > d^*$ the outer stationary point that bounds the attractive basin. The expected number of bound neighbors around one channel is

$$n_b(\sigma,\bar{\rho}) = \bar{\rho}\int_{d^*}^{d_b} 2\,\pi R\,\exp\left[-\beta\left(U_{\text{eff}}(R;\sigma) - U_{\text{eff}}(d_b;\sigma)\right)\right]\mathrm{d}R, \tag{S15}$$

where $\bar{\rho}$ is the mean areal density and $\beta = (k_B T)^{-1}$. When the attractive basin is narrow on the scale over which the Boltzmann factor varies, the integral is well approximated by

$$n_b(\sigma,\bar{\rho}) \simeq \bar{\rho}\,A_b(\sigma)\,\mathrm{e}^{\beta\varepsilon_b(\sigma)}, \tag{S16}$$

with $A_b(\sigma) \simeq 2\pi d^*(\sigma)[d_b(\sigma) - d^*(\sigma)]$, $\varepsilon_b(\sigma) = U_{\text{eff}}(d_b;\sigma) - U_{\text{eff}}(d^*;\sigma)$. The onset of clustering is defined by $n_b \sim 1$, which gives

$$\bar{\rho}_c(\sigma) \simeq \frac{\mathrm{e}^{-\beta\varepsilon_b(\sigma)}}{A_b(\sigma)}. \tag{S17}$$

This relation is the main-text onset formula. A broader attractive basin or a larger barrier lowers the density required for clustering, whereas a narrower basin or smaller barrier raises it.

In practice, when the far-field barrier is very broad, it is convenient to replace the formal outer boundary $d_b$ by an effective trapping boundary $d_b^{\text{eff}} = \min(d_b, \eta\zeta)$, where $\zeta = \xi^{-1}$ with $\eta$ of order unity. The same expressions for $A_b$ and $\varepsilon_b$ are then evaluated at $d_b^{\text{eff}}$ rather than $d_b$. This regularization leaves the onset formula unchanged while restricting the bound basin to the spatial scale relevant for finite-time clustering and finite simulation boxes.

The same attraction–repulsion balance that sets $\bar{\rho}_c$ also gives a finite cluster-size distribution. In a compact-cluster approximation, the free energy of a cluster containing $n$ channels may be written as

$$F_n \simeq -\Delta\mu\,n + \gamma\sqrt{n} + \chi n^2, \tag{S18}$$

where the bulk term $-\Delta\mu n$ represents the near-neighbor attraction, the perimeter term $\gamma\sqrt{n}$ counts channels at the boundary with fewer favorable contacts, and the quadratic term $\chi n^2$ represents the accumulated long-range repulsive overlap inside larger assemblies. The probability of observing a cluster of size $n$ is then

$$P_n \propto \exp[-\beta F_n]. \tag{S19}$$

Because $F_n$ decreases initially and then rises again, $P_n$ is broad but finite rather than monotonically favoring either isolated monomers or unlimited coarsening.

If $a_0$ is the effective projected area per channel within a compact cluster and $\ell_0$ is the corresponding perimeter scale, then $A \simeq a_0 n$ and $P \simeq \ell_0 \sqrt{n}$.

The area and perimeter distributions follow by change of variables,

$$p_A(A) \propto \exp\left[-\beta F_{\frac{A}{a_0}}\right], \qquad p_P(P) \propto P \exp\left[-\beta F_{\left(\frac{P}{\ell_0}\right)^2}\right]. \tag{S20}$$

These expressions explain why the measured cluster-area and cluster-perimeter histograms in the main text are right-skewed yet finite. Small clusters are stabilized by the attractive basin, but larger clusters are progressively disfavored by boundary loss and longer-range repulsion.

**Supplementary Note 4. Literature-based channel parameters and membrane-tension inference**

This note summarizes what can already be fixed for the channel parameters from published structural, mechanical and molecular studies of Piezo1 before comparison with cell data. The goal is to define one transferable coarse-grained Piezo1 channel model before using the experimental comparisons to extract membrane-context quantities. In this strategy, the literature is used first to assign specific working values to $a_{\mathrm{ch}}$, $\Delta A$, $\alpha_c$, $\lambda$, $\mu$ and $q_0$, after which the comparisons to imaging and patch-clamp measurements mainly constrain the effective membrane tension and the relevant Piezo1 density.

We adopt the coarse-grained convention $q = 0$ for a curved closed-like reference state and $q = 1$ for a flattened open-like reference state. With that convention, the published literature is sufficient to choose one explicit working parameter set for the channel side of the model. The adopted values are collected in Supplementary Table S3. These numbers are fixed across all experimental comparisons, so that differences among cell types are attributed to membrane-context parameters rather than to redefinition of the channel itself.

The experimental comparisons in the main text then infer membrane tension by matching the cluster scale predicted by Brownian dynamics to the cluster scale measured in imaging data. For a system with experimentally constrained density $\rho_{\mathrm{exp}}$ and measured cluster area $A_{\mathrm{exp}}$, a trial membrane tension $\sigma$ determines the pair interaction $U_{\mathrm{eff}}(R;\sigma)$ and therefore the simulated organization at the observed density. Let $A_{\mathrm{sim}}(\sigma;\rho_{\mathrm{exp}})$ denote the simulated mean or median cluster area extracted with the same segmentation rule used for the experiment. The inferred membrane tension is obtained by minimizing

$$\mathcal{E}(\sigma) = \left|\log A_{\mathrm{sim}}(\sigma;\rho_{\mathrm{exp}}) - \log A_{\mathrm{exp}}\right|. \tag{S42}$$

Equivalently, $\sigma$ may be updated iteratively until the simulated cluster scale agrees with the measured one within the chosen tolerance. This is the workflow summarized in Supplementary Fig. 6. Density alone does not uniquely determine organization, so the cluster-size comparison acts as a second constraint. In this way, red blood cells, Neuro-2a cells, and HEK293T cells can be placed on the same tension-dependent organization landscape while using the same transferable channel model.

**Supplementary Note 5. Analytical origin of the cluster-area distribution and its connection to the pair potential**

The cluster-area distribution in Fig. 3i can be understood as the finite-density consequence of the same two-body interaction that produces Piezo1 association. In the main-text flat-membrane theory, the effective pair interaction may be written schematically as

$$U_{\text{eff}}(R;\sigma) = U_{\text{hc}}(R) + B(\sigma)K_0(R/\zeta) - C(\sigma)\zeta^{-1}K_1\left(\frac{R}{\zeta}\right), \qquad \zeta = \left(\frac{\kappa_{\text{eff}}}{\sigma}\right)^{\frac{1}{2}}. \tag{S21}$$

The $K_0$ term is the longer-ranged repulsive overlap of membrane-deformation fields, whereas the $K_1$ term is the short-ranged state-shape feedback attraction generated when neighboring channels flatten one another and reduce boundary mismatch. This pair potential defines the preferred bound spacing $d^*$, the outer barrier position $d_b$, the attractive-basin width $w = d_b - d^*$, the capture area $A_b \simeq 2\pi d^* w$, and the escape barrier $\epsilon_b = U_{\text{eff}}(d_b) - U_{\text{eff}}(d^*)$. These pair-level quantities can be coarse-grained into a cluster-area free energy.

For a compact cluster of area $A$, we write

$$F(A) = -\Delta f\, A + 2\sqrt{\pi}\,\gamma A^{\frac{1}{2}} + E_{\text{rep}}(A). \tag{S22}$$

The first term is the bulk cohesive gain from short-range attractive contacts, the second is the perimeter cost of boundary channels with fewer attractive neighbors, and the third is the accumulated nonlocal repulsive energy from the repulsive part of the pair potential. The preferred spacing fixes the intracluster density,

$$\rho_{\text{in}} \simeq \frac{2}{\sqrt{3}\,d^{*2}}, \qquad N \simeq \rho_{\text{in}} A. \tag{S23}$$

The bulk drive can be related to the pair-level clustering threshold. If

$$\bar{\rho}_c \simeq \frac{\exp(-\beta\epsilon_b)}{A_b}, \tag{S24}$$

then a useful reduced form is

$$\Delta f \simeq \rho_{\text{in}} k_B T \ln\left(\frac{\bar{\rho}}{\bar{\rho}_c}\right) = \rho_{\text{in}}[\epsilon_b + k_B T \ln(\bar{\rho} A_b)]. \tag{S25}$$

Thus $\epsilon_b$ controls how strongly a channel is retained in the attractive basin, $A_b$ controls the configurational size of that basin, and $\bar{\rho}$ controls the supply of channels. The line tension is the energetic penalty for missing attractive contacts at the cluster boundary. A simple estimate is $\gamma \simeq c_\gamma \frac{\epsilon_{\text{bond}}}{d^*}$, where $c_\gamma$ is an order-one geometrical factor. In a narrow-well approximation one may take $\epsilon_{\text{bond}} \sim \epsilon_b$. More generally, $\epsilon_{\text{bond}}$ should be obtained from the full attractive-basin partition integral of $U_{\text{eff}}$, because $\epsilon_b$ is strictly an escape barrier rather than a thermodynamic bond free energy.

The repulsive cluster energy is obtained by summing the repulsive $K_0$ tail over all channel pairs in the cluster. In a continuum approximation,

$$E_{\text{rep}}(A) = \frac{\rho_{\text{in}}^2}{2}\int_{\Omega_A} d^2 r \int_{\Omega_A} d^2 r'\, U_{\text{rep}}(|\mathbf{r} - \mathbf{r}'|), \tag{S26}$$

with $U_{\text{rep}}(R) = B(\sigma)K_0(R/\zeta)\Theta(R - r_0)$.

Here $\Omega_A$ is the cluster domain and $r_0$ is a short-distance cutoff of order $d^*$ or $d_b$. The cutoff removes the near-neighbor attractive basin that has already been absorbed into the cohesive term in Eq. S22. For a circular cluster with radius $R_{\mathrm{cl}} = (A/\pi)^{1/2}$,

$$E_{\mathrm{rep}}(A) = \pi\rho_{\mathrm{in}}^2 B(\sigma) \int_{r_0}^{2R_{\mathrm{cl}}} d\, s\, s\, K_0(s/\zeta) A_{\mathrm{ov}}(s; R_{\mathrm{cl}}), \tag{S27}$$

where

$$A_{\mathrm{ov}}(s; R_{\mathrm{cl}}) = 2R_{\mathrm{cl}}^2 \cos^{-1}\left(\frac{s}{2R_{\mathrm{cl}}}\right) - \frac{s}{2}\left(4R_{\mathrm{cl}}^2 - s^2\right)^{\frac{1}{2}} \tag{S28}$$

is the overlap area of two disks of radius $R_{\mathrm{cl}}$ separated by $s$.

This expression also gives the coefficients in the large-area expansion. For $R_{\mathrm{cl}} \gg \zeta$, the overlap area has the small-$s/R_{\mathrm{cl}}$ expansion

$$A_{\mathrm{ov}}(s; R_{\mathrm{cl}}) = A - \frac{P}{\pi} s + O\left(\frac{s^3}{R_{\mathrm{cl}}}\right), \qquad P = 2\sqrt{\pi A}, \tag{S29}$$

so that

$$E_{\mathrm{rep}}(A) = e_{\mathrm{rep}} A - \gamma_{\mathrm{rep}} A^{\frac{1}{2}} + O(1). \tag{S30}$$

For the screened repulsive potential above, the bulk repulsive energy density is

$$e_{\mathrm{rep}} = \pi\rho_{\mathrm{in}}^2 B(\sigma) \int_{r_0}^{\infty} d\, s\, s K_0(s/\zeta) = \pi\rho_{\mathrm{in}}^2 B(\sigma) \zeta^2 x_0 K_1(x_0), \qquad x_0 = \frac{r_0}{\zeta}. \tag{S31}$$

The boundary correction is

$$\gamma_{\mathrm{rep}} = 2\sqrt{\pi}\rho_{\mathrm{in}}^2 B(\sigma) \int_{r_0}^{\infty} d\, s\, s^2 K_0(s/\zeta) = 2\sqrt{\pi}\rho_{\mathrm{in}}^2 B(\sigma) \zeta^3 \int_{x_0}^{\infty} d\, x\, x^2 K_0(x). \tag{S32}$$

When $r_0 \ll \zeta$, these reduce to the useful estimates

$$e_{\mathrm{rep}} \simeq \pi\rho_{\mathrm{in}}^2 B(\sigma)\zeta^2, \qquad \gamma_{\mathrm{rep}} \simeq \pi^{\frac{3}{2}}\rho_{\mathrm{in}}^2 B(\sigma)\zeta^3. \tag{S33}$$

The minus sign in Eq. S30 has a simple physical origin: channels at the cluster edge have fewer repulsive neighbors than channels in the interior, so the finite boundary reduces the total repulsive energy relative to a bulk cluster of the same area. Thus the repulsive tail contributes both a positive bulk penalty $e_{\mathrm{rep}}A$ and an effective negative line contribution $-\gamma_{\mathrm{rep}}A^{1/2}$.

Combining Eqs. S22 and S33, the far large-area free energy is

$$F(A) \simeq \left(e_{\mathrm{rep}} - \Delta f\right)A + \left(2\sqrt{\pi}\gamma - \gamma_{\mathrm{rep}}\right)A^{\frac{1}{2}} + O(1). \tag{S34}$$

Therefore the large-area tail is stable only when $e_{\mathrm{rep}} > \Delta f$. If this condition holds, very large clusters are exponentially suppressed. If $\Delta f$ exceeds $e_{\mathrm{rep}}$, the large-area free energy decreases with area and the theory predicts continued coarsening rather than a self-limited cluster distribution.

Finally, the measured area distribution is not simply $\exp[-\beta F(A)]$, because zero-area clusters are not physical and the small-cluster phase-space measure must be included. A minimal form is

$$P(A) \propto A^{\nu} \exp[-\beta F(A)]. \tag{S35}$$

The factor $A^{\nu}$ makes $P(A)$ vanish as $A \to 0$. The maximum occurs at a finite area where the short-range cohesive gain and reduction of boundary cost are balanced by the growing accumulated repulsive deformation-field energy. The large-area tail is then controlled by the rise of $E_{\mathrm{rep}}(A)$, with the far asymptotic decay set primarily by $e_{\mathrm{rep}} - \Delta f$.

Perimeter distribution

A perimeter distribution follows from the same cluster free energy once a geometrical relation between cluster area and cluster perimeter is specified. For compact, approximately circular clusters, the perimeter $L$ and area $A$ obey $L \simeq 2\sqrt{\pi A},\ A = \frac{L^2}{4\pi}$. The probability density for perimeter is then obtained by the change of variables

$$P_L(L) = P_A\left(\frac{L^2}{4\pi}\right)\frac{dA}{dL} = P_A\left(\frac{L^2}{4\pi}\right)\frac{L}{2\pi}. \tag{S36}$$

Using $P_A(A) \propto A^{\nu} \exp[-\beta F(A)]$ and Eq. S22, this gives

$$P_L(L) \propto L^{2\nu+1} \exp\left\{-\beta\left[-\frac{\Delta f}{4\pi}L^2 + \gamma L + E_{\mathrm{rep}}\left(\frac{L^2}{4\pi}\right)\right]\right\}. \tag{S37}$$

Thus, the perimeter histogram is not an independent distribution: in the compact-cluster limit it is the area distribution expressed in perimeter variables. The factor $L^{2\nu+1}$ makes the perimeter distribution vanish at $L = 0$, the maximum occurs at a finite perimeter set by the same attraction-repulsion balance that sets the preferred area, and the large-$L$ tail is suppressed by the same accumulated repulsive membrane-deformation energy.

The large-perimeter asymptotic form follows directly from Eq. S30. Substituting $A = L^2/(4\pi)$ gives

$$F_L(L) \equiv F\left(\frac{L^2}{4\pi}\right) \simeq \frac{e_{\mathrm{rep}} - \Delta f}{4\pi}L^2 + \left(\gamma - \frac{\gamma_{\mathrm{rep}}}{2\sqrt{\pi}}\right)L + O(1). \tag{S38}$$

Therefore, when $e_{\mathrm{rep}} > \Delta f$, the far perimeter tail is approximately

$$P_L(L) \sim L^{2\nu+1} \exp\left[-\beta\frac{e_{\mathrm{rep}} - \Delta f}{4\pi}L^2 - \beta\left(\gamma - \frac{\gamma_{\mathrm{rep}}}{2\sqrt{\pi}}\right)L\right]. \tag{S39}$$

The dominant quadratic term is simply the large-area exponential tail written in terms of $L^2$. The linear term contains two competing boundary effects: the attractive line tension $\gamma$, which penalizes cluster boundary, and the repulsive boundary correction $\gamma_{\mathrm{rep}}/(2\sqrt{\pi})$, which lowers the repulsive energy because edge channels have fewer repulsive neighbors.

For experimentally segmented clusters, the circular relation is only an approximation. A more general description introduces the isoperimetric shape factor

$$\chi = \frac{L^2}{4\pi A} \geq 1, \qquad A = \frac{L^2}{4\pi\chi}. \tag{S40}$$

Here $\chi = 1$ corresponds to a circular cluster, whereas $\chi > 1$ represents elongated, rough, or jagged clusters. The perimeter distribution then becomes

$$P_L(L) = \int_1^\infty d\chi\, P_A\left(\frac{L^2}{4\pi\chi}\right) P(\chi|L) \frac{L}{2\pi\chi}. \tag{S41}$$

The function $P(\chi|L)$ accounts for boundary roughness, anisotropy, localization noise, and the Voronoi/DBSCAN segmentation rule. These effects broaden the perimeter histogram more strongly than the area histogram, because perimeter is more sensitive to boundary irregularity. Nevertheless, the peak and the tail remain controlled by the same physical balance: short-range cohesive attraction favors growth, while accumulated membrane-mediated repulsion suppresses overly large clusters.

**Supplementary Note 6. Estimation of membrane tension under hypo-osmotic stimulation**

Because extracellular osmolarity does not uniquely determine membrane tension, we used representative effective membrane tensions guided by published tether-force measurements. Resting plasma-membrane tensions span approximately 0.04–0.45 pN nm$^{-1}$ across cell types. We therefore selected 0.2 pN nm$^{-1}$ as a representative baseline tension under isotonic conditions. Roffay et al. measured the acute membrane-mechanical response to hypo-osmotic stimulation and observed an approximately 1.7-fold increase in membrane tether force at 120 mOsm[1]. Assuming the effective bending rigidity is approximately unchanged between conditions, tether force scales with the square root of effective membrane tension. The corresponding fold change in tension is $\frac{\sigma_{hypo}}{\sigma_{iso}} = \left(\frac{F_{hypo}}{F_{iso}}\right)^2 \approx (1.7)^2 \approx 2.9$. Applying this approximately threefold increase to the baseline value gives $\sigma_{120\ \mathrm{mOsm}} \approx 2.9 \times 0.2 \approx 0.58 \approx 0.6\ \mathrm{pN\ nm^{-1}}$. We therefore used 0.2 pN nm$^{-1}$ for the isotonic condition and 0.6 pN nm$^{-1}$ for 120-mOsm hypo-osmotic stimulation. These values represent effective tensions chosen to capture the experimentally observed increase in membrane stress, rather than a direct conversion from osmolarity to membrane tension.

**Supplementary Note 7. Recruitment of Piezo1 to receptor-associated membrane domains**

The uniform-membrane theory developed above determines whether a Piezo1 population at a given density and far-field membrane tension lies below or above the finite-clustering threshold. We next consider a spatially heterogeneous membrane in which receptor organization or associated membrane remodeling locally changes the mechanical environment sensed by Piezo1. Rather than introducing a new receptor-Piezo1 binding interaction, we represent these heterogeneous contributions through a prescribed local mechanical field that biases the same single-channel free energy derived in Supplementary Note 1.

Let $\mathbf{R}$ denote the in-plane position of a Piezo1 channel and $\mathbf{x}_R$ the center of a receptor-associated remodeled membrane domain. The local mechanical field is written as

$$\sigma_R(\mathbf{R}) = \sigma_0 + \delta\sigma_R\, g\left(\frac{|\mathbf{R} - \mathbf{x}_R|}{\ell_R}\right) \tag{S43}$$

where $\sigma_0$ is the far-field membrane tension, $\delta\sigma_R$ is the amplitude of the local perturbation, and $\ell_R$ is its characteristic spatial range. The dimensionless profile satisfies $g(0) = 1$ and $g(\infty) = 0$, so that the field approaches $\sigma_0 + \delta\sigma_R$ at the center of the remodeled region and returns to $\sigma_0$ in the far field. In the Brownian-dynamics simulations, we use a Gaussian profile,

$$g\left(\frac{|\mathbf{R}-\mathbf{x}_R|}{\ell_R}\right) = \exp\left[-\frac{|\mathbf{R}-\mathbf{x}_R|^2}{2\ell_R^2}\right] \tag{S44}$$

The field $\sigma_R(\mathbf{R})$ should be interpreted as an effective local mechanical bias conjugate to the projected Piezo1 area. It therefore coarse-grains receptor-associated changes in membrane tension, lipid organization, curvature, cortical coupling, or protein crowding without assigning the recruitment effect to a single microscopic source.

We couple this field to the same membrane-coupled Piezo1 conformational coordinate $q$ used in the uniform-membrane theory. For the localized-field calculation, the edge-loading stiffness is evaluated at the far-field membrane state, $S_0 \equiv S_{\mathrm{mem}}(\sigma_0)$, so that the receptor-associated perturbation acts only through the local mechanical work on the Piezo1 projected area. The one-body free energy is therefore

$$f_{RP}(q,\mathbf{R}) = \frac{\mu}{2}(q-q_0)^2 - \sigma_R(\mathbf{R})\Delta A\, q + \frac{S_0}{2}\{\max[\alpha_c - \lambda q, 0]\}^2 \tag{S45}$$

This form preserves the same intrinsic channel elasticity and state-dependent boundary mismatch used for the isolated Piezo1 channel. The localized membrane field changes the preferred channel state but does not introduce an additional pairwise attraction between the receptor and Piezo1.

For the parameter range used in the receptor-associated recruitment simulations, the channel remains on the active branch, $\alpha_c - \lambda q > 0$. Equation S45 can then be written as

$$f_{RP}(q,\mathbf{R}) = \frac{1}{2}(\mu + \lambda^2 S_0)q^2 - [\mu q_0 + \sigma_R(\mathbf{R})\Delta A + \lambda S_0 \alpha_c]q + \frac{\mu q_0^2}{2} + \frac{S_0\alpha_c^2}{2} \tag{S46}$$

Minimization with respect to $q$ gives the locally preferred Piezo1 state

$$q_R^\star(\mathbf{R}) = \frac{\mu q_0 + \sigma_R(\mathbf{R})\Delta A + \lambda S_0 \alpha_c}{\mu + \lambda^2 S_0} \tag{S47}$$

If this unconstrained optimum reaches the saturation value $q_{\mathrm{sat}} = \alpha_c/\lambda$, the full clipped form in Eq. S45 should instead be minimized piecewise, as described for the isolated channel in Supplementary Note 1.

Defining $D = \mu + \lambda^2 S_0$ and $A(\sigma) = \mu q_0 + \sigma \Delta A + \lambda S_0 \alpha_c$, the minimized one-body free energy on the active branch is

$$f_1^{\,i}(\sigma) = \frac{\mu q_0^2}{2} + \frac{S_0 \alpha_c^2}{2} - \frac{A(\sigma)^2}{2D} \tag{S48}$$

The effective receptor-associated recruitment potential is defined relative to the far-field membrane state,

$$U_{RP}(\mathbf{R}) = f_1^{\,i}[\sigma_R(\mathbf{R})] - f_1^{\,i}(\sigma_0) \tag{S49}$$

This choice gives $U_{RP} \to 0$ far from the remodeled domain. A region in which $U_{RP} < 0$ lowers the free energy of a Piezo1 channel and therefore acts as a local recruitment region. A steric hard-core term $U_{\mathrm{hc}}^{RP}$ can additionally be included if the receptor-associated structure is physically inaccessible to Piezo1; it is not required for the localized recruitment field itself.

For a weak mechanical perturbation, Eq. S49 can be expanded about the far-field state,

$$U_{RP}(\mathbf{R}) \simeq -\chi_\sigma(\sigma_0)[\sigma_R(\mathbf{R}) - \sigma_0] \tag{S50}$$

where $\chi_\sigma(\sigma_0) = -\frac{\partial f_1^i}{\partial \sigma}\Big|_{\sigma_0} = \Delta A\, q_\infty(\sigma_0)$ is the mechanical susceptibility of the isolated Piezo1 channel in the active-branch approximation. Thus, the same mechanical susceptibility that allows membrane tension to shift the Piezo1 conformation also determines how strongly a localized membrane perturbation recruits the channel.

The corresponding one-body force used in the Brownian-dynamics simulations follows from

$$\mathbf{F}_{RP}(\mathbf{R}) = -\nabla U_{RP}(\mathbf{R}) = \Delta A\, q_R^\star(\mathbf{R})\nabla\sigma_R(\mathbf{R}) \tag{S51}$$

For the Gaussian field in Eq. S44,

$$\nabla\sigma_R(\mathbf{R}) = -\frac{\delta\sigma_R}{\ell_R^2}\, g\left(\frac{|\mathbf{R} - \mathbf{x}_R|}{\ell_R}\right)(\mathbf{R} - \mathbf{x}_R) \tag{S52}$$

For $\delta\sigma_R > 0$, this force biases Piezo1 toward the center of the remodeled membrane region.

In the dilute pre-nucleation regime, the local channel density follows the Boltzmann distribution associated with the one-body recruitment potential, $\rho(\mathbf{R}) = \rho_\infty \exp[-\beta U_{RP}(\mathbf{R})]$ where $\rho_\infty$ is the far-field Piezo1 density and $\beta = (k_B T)^{-1}$. Thus, a negative recruitment potential produces local enrichment while leaving the far-field channel density unchanged.

The connection between receptor-associated recruitment and finite clustering follows directly from the uniform-membrane onset condition derived in Supplementary Note 3. Because the Piezo1-Piezo1 pair potential is kept fixed at the far-field membrane state $\sigma_0$, the relevant clustering threshold remains the uniform-membrane onset density $\rho_{\mathrm{on}}(\sigma_0)$. A receptor-associated region nucleates a finite cluster when $\rho(\mathbf{R}) \geq \rho_{\mathrm{on}}(\sigma_0)$, defining the local recruitment free-energy gain as $\Delta F_{\mathrm{recruit}}(\mathbf{R}) = -U_{RP}(\mathbf{R})$, giving the critical recruitment gain

$$\Delta F_{\mathrm{recruit}}^{\mathrm{critical}} = k_B T \ln\left[\frac{\rho_{\mathrm{on}}(\sigma_0)}{\rho_\infty}\right] \tag{S53}$$

Local finite clustering is therefore predicted when $\Delta F_{\mathrm{recruit}}(\mathbf{R}) \geq \Delta F_{\mathrm{recruit}}^{\mathrm{critical}}$. This expression separates the two roles of membrane mechanics in the heterogeneous system. The localized receptor-associated field determines where Piezo1 accumulates by lowering its one-body free energy, whereas the previously derived Piezo1-Piezo1 interaction determines whether the locally accumulated channels form a stable finite assembly. A population that is subcritical in the far field can therefore cross the pre-existing clustering boundary locally without any additional receptor-mediated attraction between Piezo1 channels.

Consistent with this construction, the many-body Brownian-dynamics energy is

$$E_{\mathrm{tot}} = \sum_{i<j} U_{\mathrm{eff}}\left(\left|\mathbf{R}_i - \mathbf{R}_j\right|; \sigma_0\right) + \sum_i U_{RP}\,(\mathbf{R}_i) \tag{S54}$$

The first term is the same non-monotonic Piezo1-Piezo1 interaction derived for a uniform membrane and is evaluated at the far-field membrane state. The second term is the receptor-associated one-body positional bias. Increasing the amplitude or spatial range of the localized field increases Piezo1 recruitment and can raise the local density above $\rho_{\mathrm{on}}(\sigma_0)$, producing the transition from diffuse enrichment to receptor-associated finite clustering shown in Fig. 6.

### Supplementary Figures

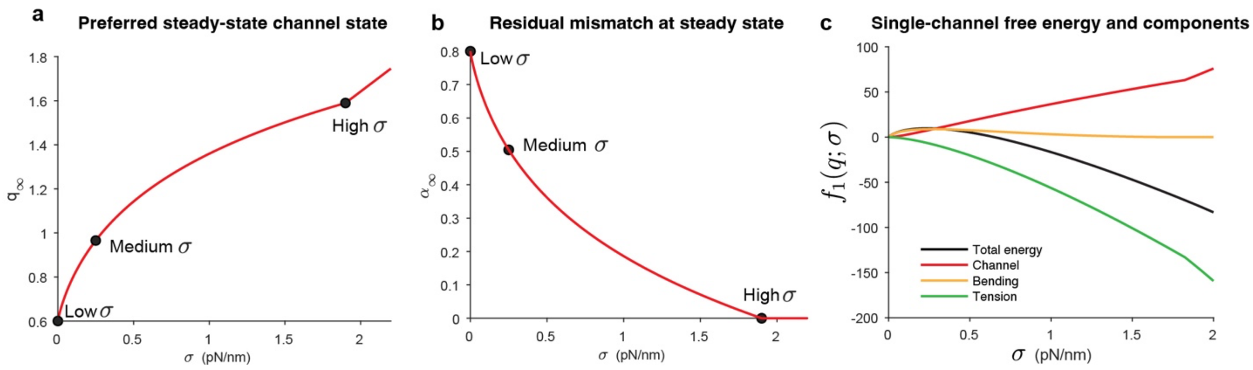


**Supplementary Figure 1: Isolated-channel mechanics determine the steady-state Piezo1 state under membrane tension. a,** Preferred steady-state channel state $q_\infty$ as a function of membrane tension $\sigma$, showing progressive flattening with increasing tension. **b,** Residual boundary mismatch $\alpha_\infty$ at steady state, which decreases as tension drives the channel toward flatter conformations. **c,** Reduced single-channel free energy and its component contributions, illustrating the balance between channel elasticity, membrane bending, and membrane tension that sets the isolated-channel state.

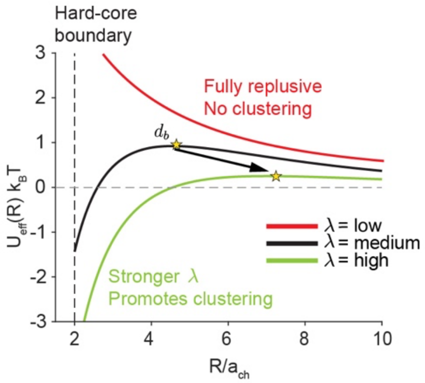


**Supplementary Figure 2: Feedback strength reshapes the Piezo1 pair potential and controls whether finite clustering can occur.** Representative pair potentials $U_{\mathrm{eff}}(R)$are shown for low, intermediate, and high feedback strength $\lambda$. Weak feedback leaves the interaction effectively repulsive, whereas stronger feedback deepens the short-range attractive basin and increases the likelihood of pair capture and finite cluster formation.

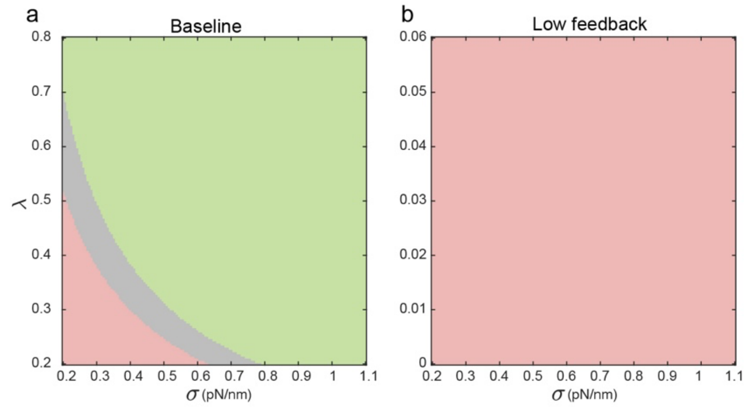


**Supplementary Figure 3: Removal of recursive feedback eliminates the finite-cluster regime. a,** Baseline phase diagram showing repulsive, hump-shaped, and fully attractive regimes when state-shape feedback is present. **b,** Low-feedback limit, in which the attractive branch is lost and the interaction collapses toward the repulsive regime, preventing clustering.

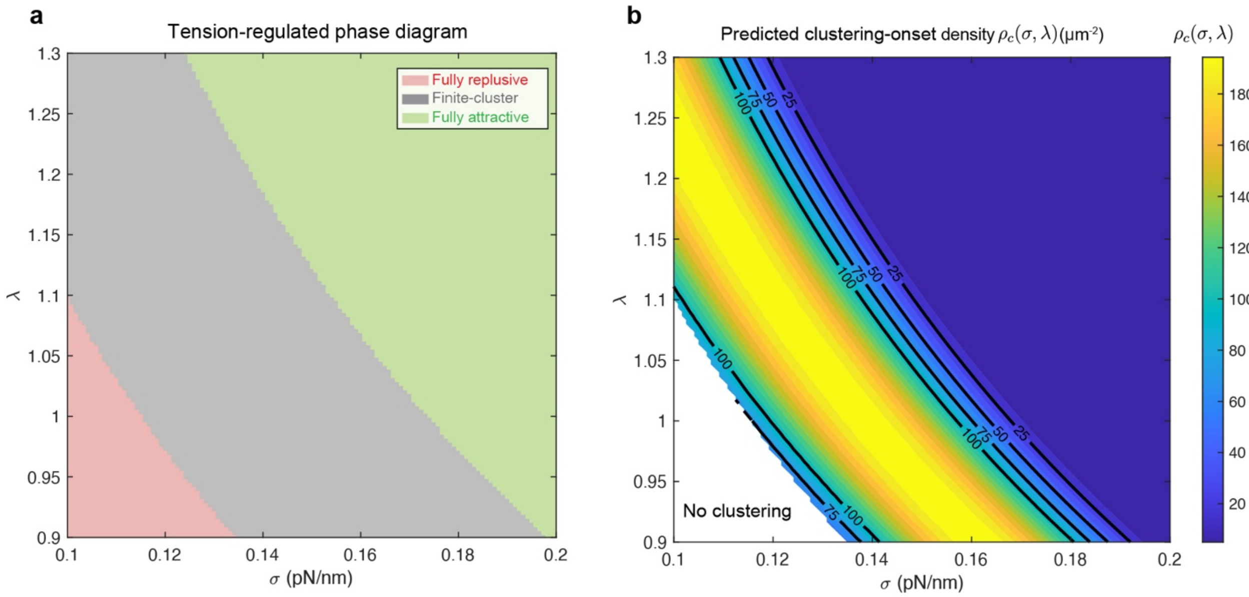


**Supplementary Figure 4: Feedback strength and membrane tension jointly determine the organization landscape and the onset density for clustering. a,** Tension-regulated phase diagram in the $(\sigma, \lambda)$plane, showing fully repulsive, hump-shaped finite-cluster, and fully attractive regimes. **b,** Predicted onset channel density $\rho_c(\sigma, \lambda)$, showing how the density threshold for clustering depends on both membrane tension and feedback strength. Contours indicate representative onset densities, and the no-clustering region corresponds to purely repulsive pair interactions.

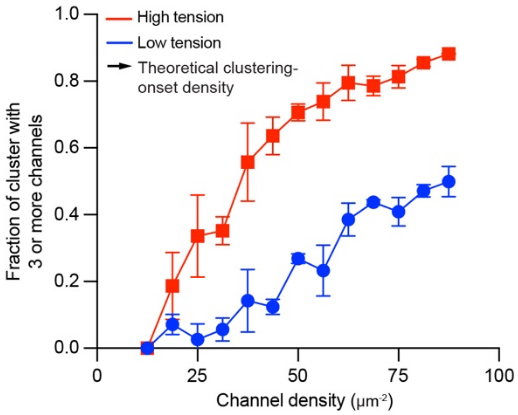


**Supplementary Figure 5: Brownian-dynamics simulations define the many-body onset of clustering.** Fraction of channels belonging to clusters of size three or larger as a function of channel density under low- and high-tension conditions. Higher membrane tension lowers the density threshold for clustering, consistent with the tension-dependent reshaping of the pair potential.

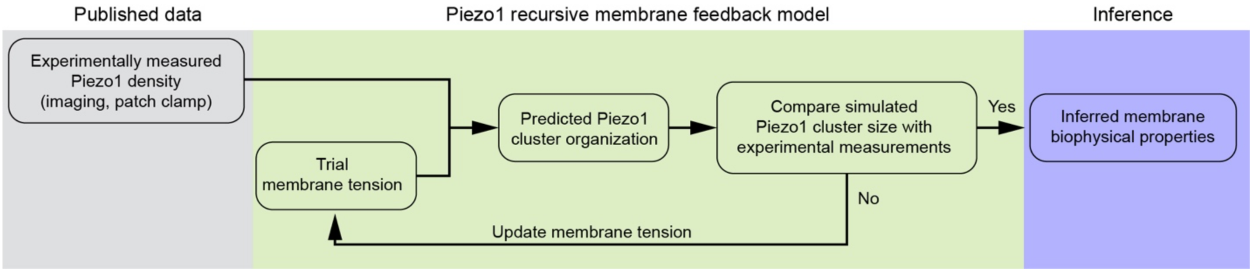


**Supplementary Figure 6: Workflow for model-constrained inference of membrane context from Piezo1 organization.** Experimentally measured Piezo1 density is used together with the recursive membrane-feedback model to generate predicted cluster organization, which is then compared with imaging-based cluster measurements. Agreement identifies the membrane-context range compatible with the observed organization, whereas disagreement is used to update the trial membrane tension iteratively.

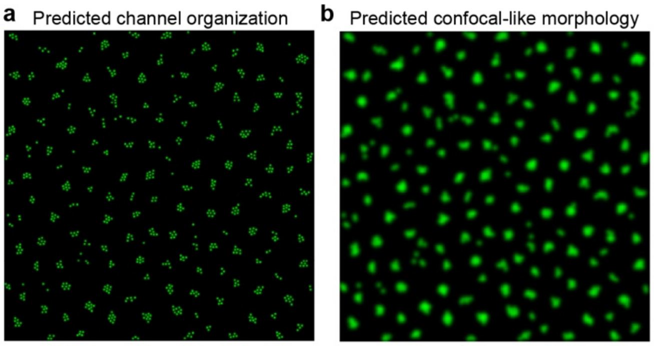


**Supplementary Figure 7: Simulated Piezo1 organization and corresponding confocal-like morphology in the finite-cluster regime. a,** Brownian-dynamics particle configuration generated from the derived pair potential. **b,** Confocal-like rendering of the same configuration after convolution with the experimental imaging kernel, illustrating how finite assemblies appear in diffraction-limited microscopy.

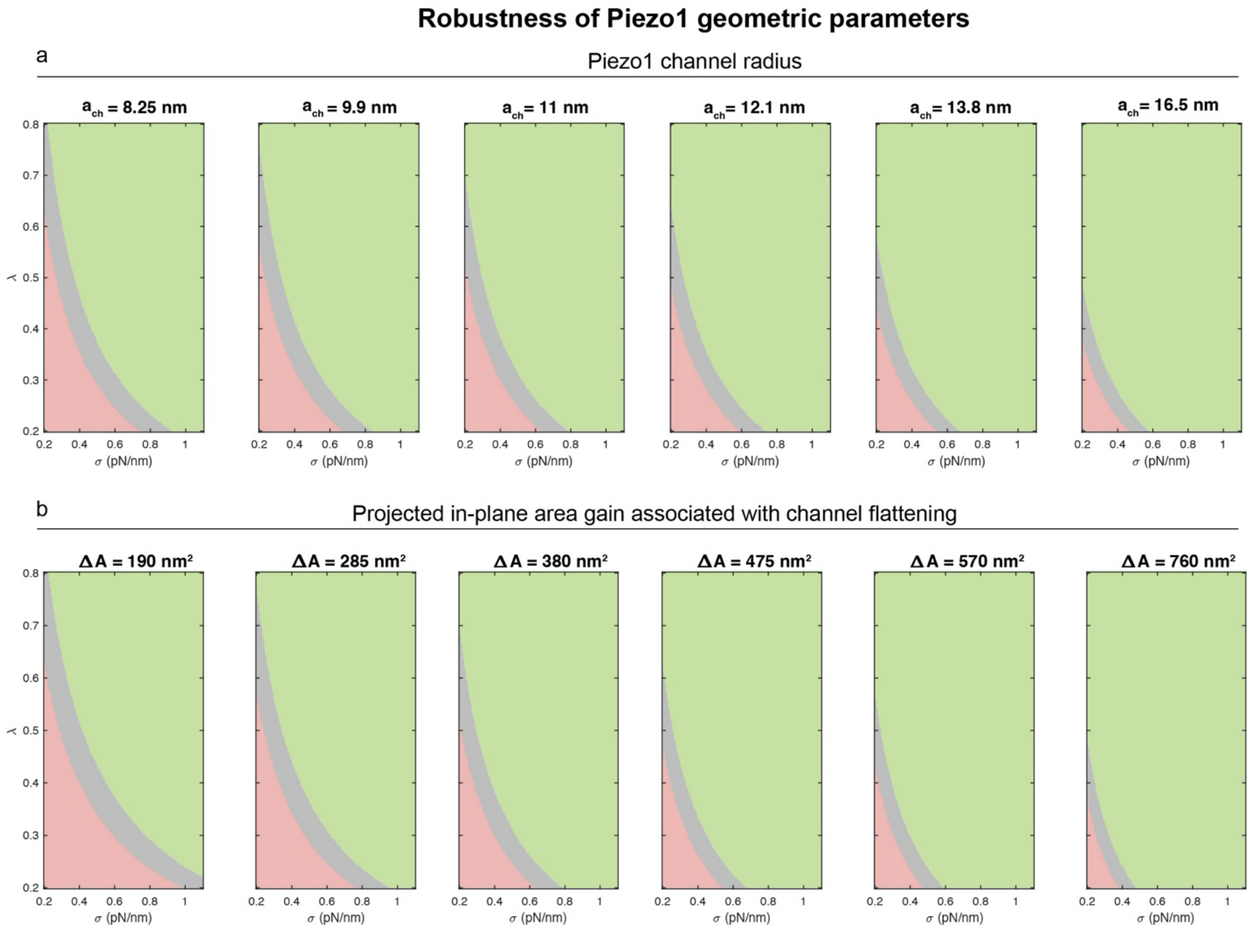


**Supplementary Figure 8: Robustness of the organization landscape to variations in geometric Piezo1 parameters.** Phase diagrams are shown while varying **a,** the effective channel radius $a_{\mathrm{ch}}$ and **b,** the projected in-plane area gain $\Delta A$ associated with flattening. Across these geometric perturbations, the finite-cluster regime persists, although its boundaries shift quantitatively.

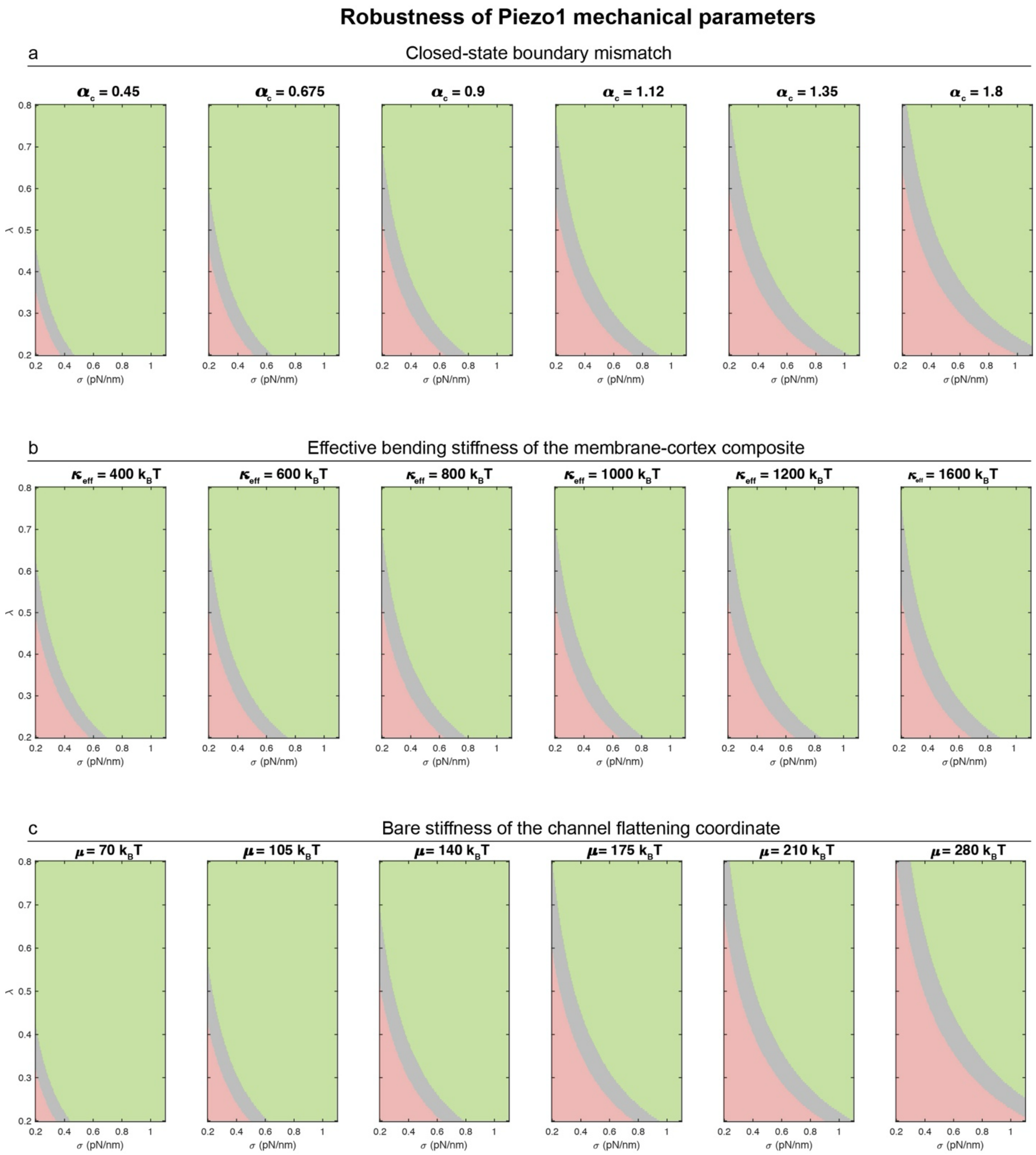


**Supplementary Figure 9: Robustness of the organization landscape to variations in mechanical Piezo1 and membrane parameters.** Phase diagrams are shown while varying **a,** the closed-state preferred boundary mismatch $\alpha_c$, **b,** the effective bending stiffness of the membrane-cortex composite $\kappa_{\text{eff}}$, and **c,** the bare stiffness $\mu$of the channel flattening coordinate. The existence of the finite-cluster regime is preserved over broad parameter ranges, indicating that the predicted organization landscape is mechanically robust.

### Supplementary Tables

#### Supplementary Table 1: Independent membrane and channel parameters appearing in the main-text Hamiltonian and boundary condition.

| Symbol | Meaning | Dimension |
|---|---|---|
| **Membrane parameters** | | |
| $\kappa_{\mathrm{eff}}$ | Effective bending stiffness of the membrane-cortex composite | $E$ |
| $\sigma$ | Membrane tension, i.e. the coefficient of the surface-gradient term $\lvert\nabla u\rvert^2$ | $EL^{-2}$ |
| **Channel parameters** | | |
| $\mu$ | Bare stiffness of the channel flattening coordinate | $E$ |
| $q_0$ | Bare preferred channel state in the absence of stress and curvature | $1$ |
| $\Delta A$ | Projected in-plane area gain associated with channel flattening; distinct from the footprint area $\pi a_{\mathrm{ch}}^2$ | $L^2$ |
| $\alpha_c$ | Closed-state preferred boundary slope mismatch | $1$ |
| $\lambda$ | Coupling coefficient in the active branch $\alpha(q) = \mathrm{pospart}(\alpha_c - \lambda q)$ that converts flattening into reduced boundary mismatch until the mismatch saturates at zero | $1$ |
| $a_{\mathrm{ch}}$ | Effective channel footprint radius at which the boundary condition is imposed | $L$ |

**Supplementary Table 2: Supplementary symbol table collecting variables, reduced coefficients, and derived observables used in the main text and Supplementary Notes.**

| Symbol | Meaning | Dimension |
|---|---|---|
| $\mathbf{r}$, $\mathbf{R}_i$ | In-plane coordinate and center position of channel $i$ | $L$ |
| $u$ | Channel-induced deformation | $L$ |
| $H_0$ | Background curvature | $L^{-1}$ |
| $q_i$ | Internal flattening coordinate of channel $i$; larger $q_i$ denotes a flatter and more open state | 1 |
| $q_{\mathrm{sat}}$ | Saturation state at which the preferred boundary mismatch reaches zero, $q_{\mathrm{sat}} = \alpha_c/\lambda$ | 1 |
| $\alpha(q_i)$ | State-dependent preferred boundary slope mismatch, $\alpha(q_i) = \mathrm{pospart}(\alpha_c - \lambda q_i)$ | 1 |
| $F$, $F^*_{\mathrm{mem}}$, $F_{\mathrm{gate}}$ | Total free energy and its membrane and gating contributions | $E$ |
| $C_i$, $ds$, $\partial_n$ | Channel boundary, boundary line element, and outward normal derivative | $L$, $L$, 1 |
| $z$ | Dimensionless footprint-tension parameters, $z = \xi a_{\mathrm{ch}}$ | 1, 1 |
| $S_{\mathrm{mem}}$ | Effective edge-loading stiffness after elimination of the membrane deformation field | $E$ |
| $f_1(q;\sigma)$ | Reduced one-body free energy of an isolated channel on a flat background | $E$ |
| $\mu_{\mathrm{eff}}$ | Renormalized stiffness after eliminating the membrane field | $E$ |
| $q_\infty$, $\alpha_\infty$ | Preferred isolated-channel state and its corresponding residual mismatch | 1, 1 |
| $R$ | Center-to-center channel separation or receptor-channel distance, depending on context | $L$ |
| $U_{\mathrm{hc}}(R)$ | Hard-core repulsion between channel footprints | $E$ |
| $J(R)$, $G(R)$ | Direct-overlap and feedback kernels in the pair interaction | $E$, $E$ |
| $J_0$, $G_0$ | Kernel amplitudes in the channel-pair interaction | $E$, $EL$ |
| $K_0$, $K_1$ | Modified Bessel functions of the second kind | 1 |
| $\xi$ | Inverse screening length, $\xi = \sqrt{\sigma/\kappa_{\mathrm{eff}}}$ | $L^{-1}$ |
| $U_{\mathrm{eff}}(R;\sigma)$ | Effective pair interaction after eliminating the channel state | $E$ |
| $B(\sigma)$, $C(\sigma)$ | Amplitudes of the repulsive $K_0$ and attractive $\xi K_1$ terms in $U_{\mathrm{eff}}$ | $E$, $EL$ |
| $\zeta$ | Screening length, $\zeta = \xi^{-1} = \sqrt{\kappa_{\mathrm{eff}}/\sigma}$ | $L$ |
| $d^*$, $d_b$, $w$ | Preferred bound spacing, outer barrier radius, and outward basin width $w = d_b - d^*$ | $L$, $L$, $L$ |
| $A_b$ | Narrow-well capture area, $A_b \simeq 2\pi d^* w$ | $L^2$ |
| $\varepsilon_b$ | Escape barrier from the preferred bound state, $\varepsilon_b = U_{\mathrm{eff}}(d_b) - U_{\mathrm{eff}}(d^*)$ | $E$ |
| $\bar{\rho}$, $\bar{\rho}_c$ | Mean areal density and clustering threshold density | $L^{-2}$ |
| $\beta$, $k_B T$ | Inverse thermal energy and thermal energy | $E^{-1}$, $E$ |
| $F_n$ | Effective free energy of a cluster containing $n$ channels | $E$ |
| $\Delta\mu$ | Bulk free-energy gain per channel upon entering a cluster | $E$ |
| $\gamma$ | Effective boundary free-energy coefficient for a compact cluster | $E$ |
| $\chi$ | One-body response coefficient, $\chi = a_{\mathrm{ch}}\lambda S_{\mathrm{mem}}$ | $EL$ |
| $a_0$, $\ell_0$ | Effective projected area per channel and perimeter scale in compact-cluster geometry | $L^2$, $L$ |
| $U^{\mathrm{RP}}_{\mathrm{hc}}(R)$ | Hard-core exclusion between a receptor and a Piezo1 channel | $E$ |
| $U_{\mathrm{RP}}(R)$ | Effective defect-channel interaction | $E$ |
| $\rho(R)$, $\rho_\infty$ | Radial channel density around a defect and its far-field value | $L^{-2}$, $L^{-2}$ |

**Supplementary Table 3: Adopted literature-based values for the independent Piezo1 channel parameters in Supplementary Table 1.**

| Parameter | Adopted value | Literature basis | Role in the present coarse-grained model |
|---|---|---|---|
| $a_{\mathrm{ch}}$ | 11 nm | Structural and footprint studies place the relevant Piezo1 boundary scale in the $\sim 10$–$14$ nm range, depending on whether one tracks the protein core or the membrane-supported nanodome footprint[2-4]. We adopt $11$ nm as the boundary-condition radius of the coarse-grained channel. | Sets the channel length scale entering the boundary condition and fixes the near-contact preferred spacing scale $2a_{\mathrm{ch}} \simeq 22$ nm. |
| $\Delta A$ | $300$ nm$^2$ | The curved-to-flattened lipid-membrane structure of Piezo1 shows an in-plane area expansion of approximately $300$ nm$^2$ [5]. | Sets the tension work term $-\sigma \Delta A\, q$ and therefore the direct stress sensitivity of the coarse-grained channel state. |
| $\alpha_c$ | $0.8$ | Structural analyses of the resting Piezo dome imply a dome-membrane contact angle of order $40°$[6]. Taking the boundary mismatch as the corresponding slope gives $\alpha_c \sim \tan 40° \approx 0.84$, for which we adopt the rounded coarse-grained value $0.8$. | Closed-state preferred boundary mismatch. This fixes the amplitude of the elastic field produced by an isolated curved channel. |
| $\lambda$ | $0.81$ | Taking $\alpha_c \simeq 0.8$ together with $\mu \simeq 140\, k_B T$, $\Delta A \simeq 300$ nm$^2$, and the reported single-channel opening scale $\sigma_{1/2} \simeq 1.9$ pN nm$^{-1}$ [5] implies that the active branch should be driven close to zero mismatch by opening-scale flattening, which gives $\lambda$ of order $0.8$. Crowding simulations independently support the sign of this feedback because neighbor-induced footprint overlap flattens the channel and promotes opening[7]. | Encodes recursive state-shape feedback. With this choice the residual mismatch is driven to zero around the single-channel opening scale rather than becoming negative at larger flattening. |
| $\mu$ | $140\, k_B T$ | The structural lipid-membrane study reports a half-maximal activation tension of about $1.9$ pN nm$^{-1}$ together with $\Delta A \simeq 300$ nm$^2$[5]. Their product gives an energy scale $\sigma_{1/2}\Delta A \simeq 570$ pN nm $\simeq 140\, k_B T$, which we adopt as the coarse-grained flattening stiffness. This opening scale corresponds to a membrane tension of order $1$–$2$ pN nm$^{-1}$ and, for an effective mechanically coupled membrane-cortex thickness of $\sim 0.2\,\mu$m, to a cell-scale stress of order $10$ kPa. | Bare stiffness of the internal state coordinate after normalizing the full curved-to-flat conformational excursion to $q \in [0,1]$. |
| $q_0$ | $0$ | By construction the zero-stress reference state is chosen as the curved closed-like state, consistent with the resting curved structures[1,2,5,8-10]. | Places the bare minimum of the coarse-grained state variable at the curved end of the interval before membrane stress or crowding shifts it. |
